# ISG15 drives immune pathology and respiratory failure during viral infection

**DOI:** 10.1101/2020.04.13.039321

**Authors:** Namir Shaabani, Jaroslav Zak, Jennifer L. Johnson, Zhe Huang, Nhan Nguyen, Daniel C. Lazar, Vincent F. Vartabedian, Nadine Honke, Marco Prinz, Klaus-Peter Knobeloch, Kei-ichiro Arimoto, Dong-Er Zhang, Sergio D. Catz, John R. Teijaro

## Abstract

Cytokine storm during respiratory viral infection is an indicator of disease severity and poor prognosis. Type 1 interferon (IFN-I) production and signaling has been reported to be causal in cytokine storm-associated pathology in several respiratory viral infections, however, the mechanisms by which IFN-I promotes disease pathogenesis remain poorly understood. Here, using *Usp18-*deficient, USP18 enzymatic-inactive and *Isg15*-deficient mouse models, we report that lack of deISGylation during persistent viral infection leads to severe immune pathology characterized by hematological disruptions, cytokine amplification, lung vascular leakage and death. This pathology requires T cells but not T cell-intrinsic deletion of *Usp18*. However, lack of *Usp18* in myeloid cells mimicked the pathological manifestations observed in *Usp18*^*-/-*^ or *Usp18*^C61A^ mice which were dependent on *Isg15*. We further mechanistically demonstrate that interrupting the ISGylation/deISGylation circuit increases extracellular levels of ISG15 which is accompanied by inflammatory neutrophil accumulation to the lung. Importantly, neutrophil depletion reversed morbidity and mortality in *Usp18*^C61A^ mice. In summary, we reveal that the enzymatic function of *Usp18* is crucial for regulating extracellular release of ISG15. This is accompanied by altered neutrophil differentiation, cytokine amplification and mortality following persistent viral infection. Moreover, our results suggest that extracellular ISG15 may drive the inflammatory pathology observed and could be both a prospective predictor of disease outcome and a therapeutic target during severe respiratory viral infections.

## Introduction

Type I interferon (IFN-I) is one of the first cytokines induced during viral infection and plays an essential role in orchestrating effective immune responses due to its induction of antiviral genes and as an immune stimulant promoting cytokine/chemokine production, upregulating activation receptors, recruiting immune cells to infected tissues and facilitating optimal antigen presentation to T cells (McNab et al., 2015). Despite the central role of IFN-I in promoting an antiviral state, deleterious consequences of IFN-I signaling during viral infection have been reported (Ng et al., 2015; Shaabani et al., 2018; Teijaro et al., 2013; Wilson et al., 2013). An elevated IFN-I gene signature correlates with disease severity following influenza virus, lymphocytic choriomeningitis virus and severe acute respiratory syndrome (SARS)-Coronavirus (SARS-CoV) infections (Baccala et al., 2014; Channappanavar et al., 2016; Davidson et al., 2014; Davidson et al., 2015; Teijaro, 2016; Teijaro et al., 2011). Moreover, genetic or pharmacological inhibition of IFN-I signaling or production alleviates pathological manifestations and mortality following Influenza and SARS-CoV infection in mice (Channappanavar et al., 2016; Davidson et al., 2014; Teijaro et al., 2011). In specific mouse models of arenavirus infection, elevated levels of IFN-I correlate with an Acute Respiratory Distress-like Syndrome (ARDLS); severe lung immune pathology, vascular leakage and death. Inhibition of IFN-I signaling reduces lung pathology and promotes survival in these models (Baccala et al., 2014; Oldstone et al., 2018; Schnell et al., 2012). Moreover, elevated expression of interferon stimulated genes (ISGs) generally correlates with the severity of ARDS pathology (Nick et al., 2016). Despite a strong link between IFN-I signaling and immune pathology during viral infection, the role and mechanisms by which specific downstream ISGs contribute to disease outcome is lacking.

The binding of IFN-α or -β to the interferon-α/β receptor (IFNAR) complex induces the expression of hundreds of genes. One of the strongest interferon-induced genes is interferon-stimulated gene 15 *(Isg15*), which encodes for the ISG15 protein, the first identified ubiquitin-like modifier. Two major functions of ISG15 have been described to date; during a three-step enzymatic cascade termed ISGylation (Malakhov et al., 2002), ISG15 covalently conjugates to proteins where it can regulate protein stability, trafficking and function. In addition, ISG15 can be secreted from cells and act as a cytokine-like molecule to modulate immune cell function (Perng and Lenschow, 2018). While the role of conjugated ISG15 has been extensively studied with respect to control of viral and bacterial infections, studies investigating the known conjugation of ISG15 to both host and viral proteins have primarily focused on how ISGylation inhibits viral replication. However, the biological relevance of the balance between ISGylation and deISGylation on innate and adaptive immune cell functions remains poorly studied.

Another ISG induced following IFN-I signaling is Ubiquitin-specific protease 18 (Usp18). Functionally, USP18 inhibits IFNAR signaling in a negative feedback loop (Malakhova et al., 2006) and, via its isopeptidase enzymatic function, cleaves ISG15 from target proteins in a process called deISGylation (Malakhov et al., 2002). A cysteine-to-alanine codon mutation in USP18 protein at position 61 in mice (*Usp18*^C61A^ knock-in mice (Dauphinee et al., 2014; Ketscher et al., 2015)) or at position 64 in humans abolishes this isopeptidase activity while preserving the protein’s ability to antagonize IFN-I signaling (Ketscher et al., 2015; Malakhova et al., 2006).

Humans with recessive mutations in *USP18* display severe pathological manifestations with heightened innate inflammation, neurological abnormalities, respiratory failure and thrombocytopenia, and, as a result, all known patients have succumbed shortly after birth (Alsohime et al., 2020; Meuwissen et al., 2016). Pathological manifestations in *USP18*-deficient humans are associated with heightened IFN-I signaling, however, the contribution of dysregulated deISGylation in *USP18*-deficient patients remains unclear. Patients with null mutations in *ISG15* have also been reported and display enhanced anti-viral capacity, which is attributed to ISG15 mediated stabilization of the USP18 protein and is specific to human ISG15 (Speer et al., 2016). Interestingly, *ISG15*-null patients have a much less severe phenotype as compared to *USP18*-null patients despite destabilization of USP18 protein and significant elevations in IFN-I signaling, suggesting that lack of deISGylation and ISG15 may significantly contribute to pathogenesis in *USP18*-null patients. However, direct evidence for this is currently lacking.

Herein, we report that the enzymatic domain of USP18 is essential for preventing excessive accumulation of extracellular ISG15 during persistent viral infection. The accumulation of extracellular ISG15 in *Usp18*^*C61A*^ knock-in mice resulted in severe inflammatory lung pathology and mortality after viral infection. We report that the respiratory pathology associated with persistent viral infection is CD8 T cell dependent. Although T cell-selective *Usp18* deletion does not reproduce lung pathology, elevated cytokine production or mortality, we show that *Usp18*-deletion in LysM^+^ cells results in elevated levels of extracellular ISG15 accompanied by cytokine storm, lung pathology and mortality. Further, we demonstrate that this accumulation of extracellular ISG15 correlates with augmentation of CXCR4^+^ neutrophils in the lung and that depletion of neutrophils in *Usp18*^*C61A*^ mice following persistent viral infection inhibited lung pathology, cytokine amplification and promoted survival. Importantly, deletion of *Isg15* from *Usp18*^*C61A*^ mice reduced the accumulation of CXCR4^+^ neutrophils, lessened lung inflammatory pathology, suppressed cytokine amplification and restored survival. Our data suggest that extracellular levels of ISG15 could predict severe lung pathological responses in viral infection and humans with *USP18*-deficiency.

## Results

### *Usp18*^*-/-*^ mice display enhanced mortality following persistent LCMV infection

The role of IFN-I in controlling viral infections is complex, as both enhanced stimulation and inhibition of IFN-I signaling can accelerate viral control during acute or persistent viral infections, respectively (Ritchie et al., 2004; Teijaro et al., 2013; Wilson et al., 2013). Given the hypersensitivity of *Usp18*^*–/–*^ mice to IFN-I signaling, we were curious how these mice would respond to systemic LCMV infection. To study systemic LCMV infection, we used two viral clones: LCMV-Armstrong 53b (Armstrong) and -Clone 13 (Cl13). Infection with a high dose of Armstrong results in a robust anti-viral T cell response that clears the virus in 5-8 days while infection with a high dose of Cl13 results in a pan immune-suppressive state, T cell exhaustion and virus persistence for >60 days (Oldstone, 2013). Following infection with a high dose Cl13, we found that, instead of exhibiting enhanced resistance to infection, *Usp18*^*–/–*^ mice rapidly succumbed to the infection by day 8 (**Figure 1A**). Surprisingly, infection of *Usp18*^*–/–*^ mice with Armstrong also resulted in exacerbated mortality with all animals succumbing to the infection within a week, while control mice exhibited no signs of disease and survived (**Figure 1A**). Infection of *Usp18*^*–/–*^ mice with either Armstrong or Cl13 resulted in several physical signs of morbidity including ruffled appearance, hunched posture, labored breathing and reduced mobility, indicating severe illness. We next characterized the biological reasons for the exacerbated morbidity and mortality observed in *Usp18*^*–/–*^ mice using Cl13 infection model where IFN-I levels and signaling are elevated (Teijaro et al., 2013; Wilson et al., 2013). We detected distinct cytopenia punctuated by reductions in platelets, total white blood cells and lymphocyte counts in the blood of *Usp18*^*–/–*^ mice compared to control mice (**Figure 1B**). Additionally, histological examination of the lung revealed severe pathological manifestations in *Usp18*^*–/–*^ animals including alveolar wall thickening, mononuclear cell infiltrates and edema, directly reflecting the physical respiratory distress observed in *Usp18*^*–/–*^ mice during Cl13 infection (**Figure 1C**). Analysis of the bronchoalveolar lavage fluid (BALF) showed cytokine amplification in the BALF of *Usp18*^*–/–*^ compared to WT mice following Cl13 infection with significant elevations of IL-6, TNF-α, IFN-γ, GM-CSF and G-CSF, the chemokine MCP-1 and the neutrophil chemoattractant KC (**Supplemental Figure 1A**). Furthermore, we observed significant increases in total protein and lactate dehydrogenase (LDH) in the airways of *Usp18*^*–/–*^ mice compared to WT controls (**Supplemental Figure 1B**), indicating increased lung vascular permeability. Taken together, these data show *Usp18*^*–/–*^ mice infected with either acute or persistent strains of LCMV develop grave pathological manifestations that mirror symptoms observed in *Usp18*-deficient patients (Meuwissen et al., 2016) and during severe respiratory viral infections (Channappanavar et al., 2016; Teijaro, 2015; Walsh et al., 2011).

**Figure 1.**
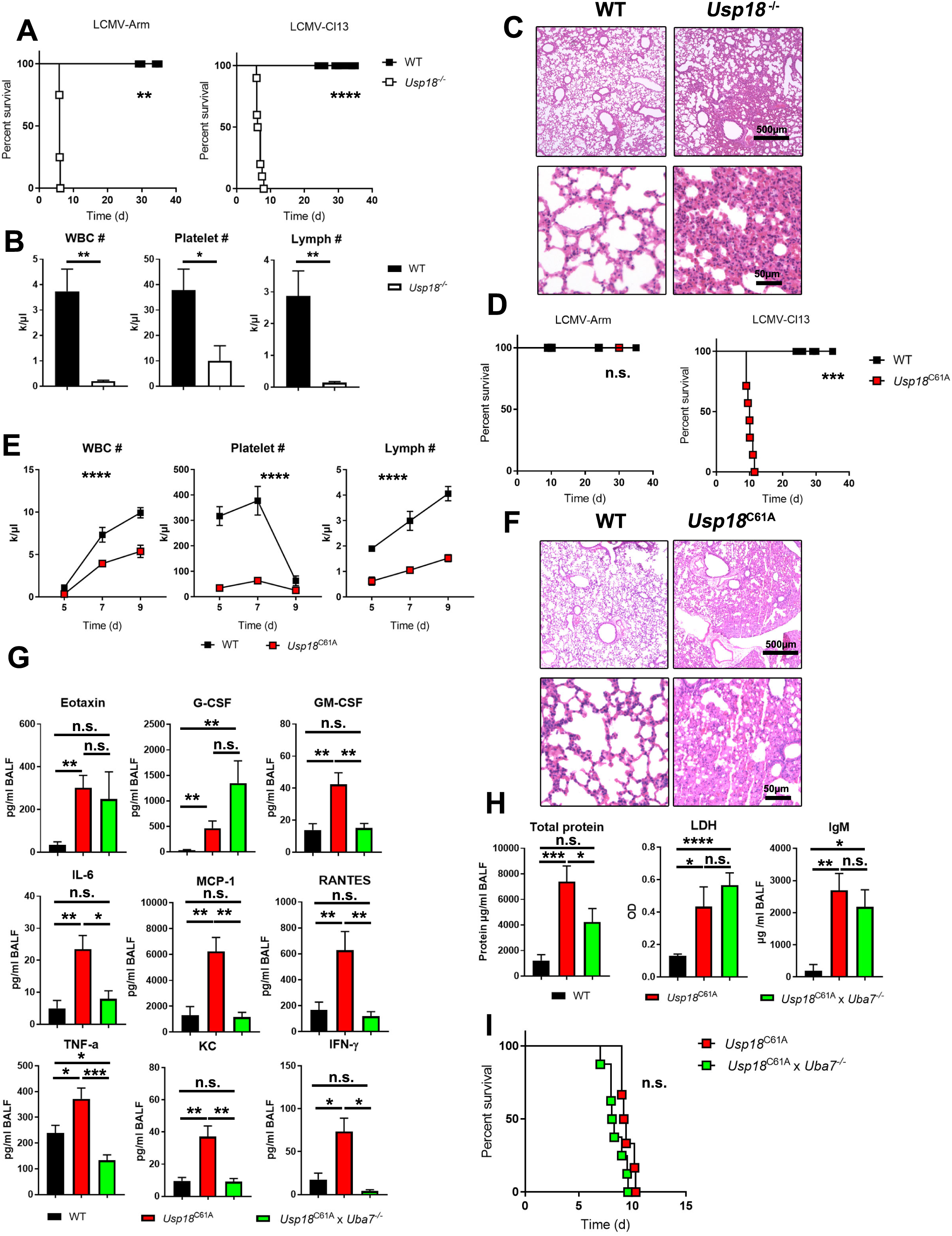
Lack of USP18 isopeptidase domain leads to lung immunopathology and acute respiratory distress-like syndrome (ARDLS). (A) WT and *Usp18*^*–/–*^ mice were infected with 2×10^6^ PFU of LCMV-Arm or 2×10^6^ PFU LCMV-Cl13. Mice survival was monitored (n = 4 mice/group Arm; 10 mice/group Cl13). (B) WT and *Usp18*^*–/–*^ mice were infected with 2×10^6^ PFU of LCMV- Cl13. On day 5, White blood cells (WBC), platelets and lymphocytes were measured in the blood (n = 7-8 mice/group). (C) WT and *Usp18*^*–/–*^ mice were infected with 2×10^6^ PFU of LCMV-Cl13. 6 days post infection, lung sections were stained with hematoxylin and eosin (n = 3 mice/group). (D) WT and *Usp18*^C61A^ mice were infected with 2×10^6^ PFU of LCMV-Arm or 2×10^6^ PFU LCMV-Cl13. Mice survival was monitored. (n= 5-7 mice/group). (E) WT and *Usp18*^C61A^ mice were infected with 2 ×10^6^ PFU LCMV-Cl13, White blood cells (WBC) platelets and lymphocytes were measured in the blood at the indicated time points (n = 5-8 mice/group). (F) Lung section of WT and *Usp18*^C61A^ mice infected with 2 ×10^6^ PFU LCMV-Cl13 for 8 days were stained with hematoxylin and eosin (n = 3 mice/group). (G-H) WT and *Usp18*^C61A^ and *Usp18*^C61A^ x *Uba7*^*-/-*^ mice were infected with 2×10^6^ PFU of LCMV-Cl13. 8 days post infection, (G) indicated cytokines (H) Total protein, LDH and IgM were analyzed in the BALF (n = 7-8 mice/group). (I) WT and *Usp18*^C61A^ and *Usp18*^C61A^ x *Uba7*^*-/-*^ mice were infected with 2×10^6^ PFU of LCMV-Cl13. Mice survival was monitored. (n= 6-8 mice/group). n.s., not significant; *P < 0.05 and **P < 0.01***P < 0.001 and ****P < 0.0001. Data are representative of at least two independent experiments

### Enzymatically inactive USP18 results in pulmonary pathology and mortality following persistent virus infection

The USP18 protein possesses two major functional domains. One inhibits IFN-I signaling through interactions with IFNAR2 and STAT2, and the other enzymatically removes ISG15 from its protein substrates in a process termed deISGylation (Arimoto et al., 2017; Malakhov et al., 2002; Malakhova et al., 2006). To mechanistically determine why *Usp18*-deficient mice exhibit severe pathological manifestations during LCMV infection, we infected WT and *Usp18*^C61A^ mice, which have normal IFN-I signaling but are unable to remove (deISGylate) ISG15 from host proteins (Ketscher et al., 2015), with Cl13 or Armstrong. Similar to *Usp18*^*–/–*^ mice, Cl13 infection of *Usp18*^C61A^ mice resulted in complete mortality (**Figure 1D**), whereas acute Armstrong infection resulted in no observable pathological manifestations (**Figure 1D**). To confirm that *Usp18*^C61A^ mice displayed normal IFN-I signaling during Cl13 infection, we measured ISGs in spleen and lung of WT and *Usp18*^C61A^ mice 2 days post-infection and found that they were induced to similar levels in WT and *Usp18*^C61A^ mice following Cl13 infection in both tissues (**Supplemental Figure 2A-B**). Further, *Usp18*^C61A^ and WT mice produced similar levels of IFN-α and β in the plasma following Cl13 infection (**Supplemental Figure 2C**). Despite normal IFN-I production and signaling, post-infection, *Usp18*^C61A^ mice exhibited pathological manifestations similar to the severe manifestations observed in *Usp18*^*–/–*^ mice including thrombocytopenia and hematological abnormalities (**Figure 1E**). Further, histological examination of the lungs of *Usp18*^*C61A*^ mice following Cl13 infection revealed severe pathological manifestations similar to those observed in *Usp18*^*–/–*^ animals including alveolar wall thickening, mononuclear cell nfiltrates and edema (**Figure 1F**). BALF cytokine level changes also mirrored those observed in *Usp18*^*–/–*^ mice following Cl13 infection, with significant elevations of IL-6, TNF-α, IFN-γ, GM-CSF, G-CSF, MCP-1 and the neutrophil chemoattractant KC observed in *Usp18*^C61A^ mice compared to WT controls (**Figure 1G**). Furthermore, we observed significant increases in total protein, lactate dehydrogenase (LDH) and IgM in the airways of *Usp18*^*C61A*^ mice compared to WT controls (**Figure 1H**), indicating increased lung vascular permeability. This enhanced pathology was not due to altered viral control, as viral titers in *Usp18*^C61A^ mice were equivalent to WT controls (**Supplementary Figure 3**). These results show that *Usp18*^C61A^ mice phenocopy *Usp18*^*–/–*^ mice post-infection and suggest that the inability to remove ISG15 from protein targets and the resultant hyper-ISGylation leads to severe lung immune pathology independent of viral loads and enhancement of IFN-I signaling.

**Figure 2.**
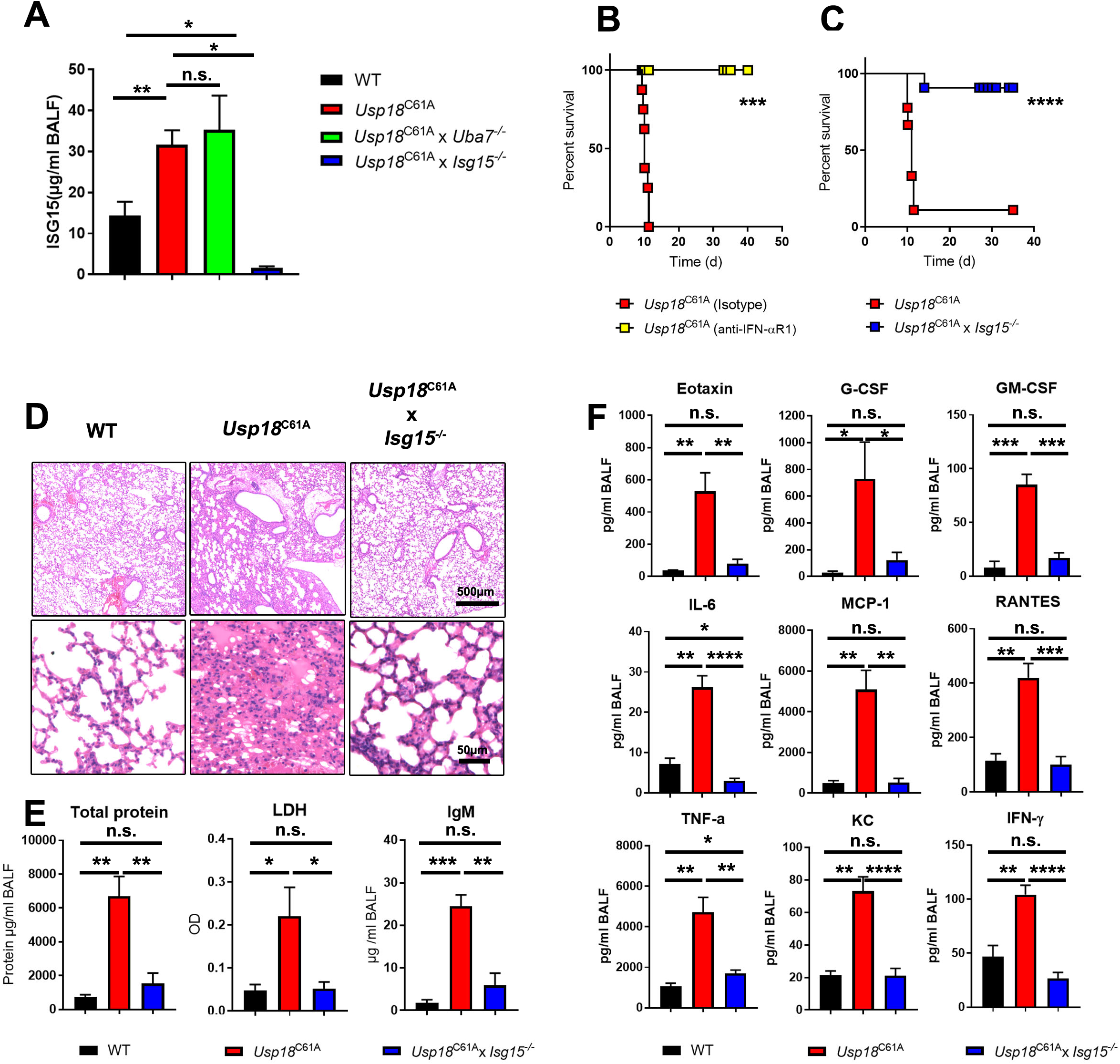
Lack of free ISG15 rescue *Usp18*^C61A^ mice during Cl3 infection. (A) WT, *Usp18*^C61A^, *Usp18*^C61A^ x *Uba7*^*-/-*^ and *Usp18*^C61A^ x *Isg15*^*–/–*^ mice were infected with 2 x10^6^ PFU LCMV-Cl13. ISG15 levels were measured in the BALFs 8 days post infection by ELISA assay (n = 4-10 mice/group) (B) WT and *Usp18*^C61A^ mice were treated with anti-IFN-αR1 antibody at day -1. Mice were infected with 2 x10^6^ PFU LCMV-Cl13 and survival was monitored (n = 8 mice/group). (C) *Usp18*^C61A^ and *Usp18*^C61A^ x *Isg15*^*–/–*^ mice were infected with 2 x10^6^ PFU LCMV-Cl13. Mice survival was monitored. (n= 9-11 mice/group). (D-F) WT, *Usp18*^C61A^ and *Usp18*^C61A^ x *Isg15*^*–/–*^ mice were infected with 2 ×10^6^ PFU LCMV-Cl13. (D) 8 days post clone-13 infection lung sections were stained with hematoxylin and eosin (n = 3 mice/group). (E) Total protein, IgM and LDH were measured by BCA or ELISA assays from BALF fluid (n = 4-5 mice/group). (F) Indicated cytokines were analyzed in the BALF by multiplex assay (n = 4-5 mice/group). n.s., not significant; *P < 0.05; **P < 0.01; ***P < 0.001 and ****P < 0.0001. Data are representative of at least two independent experiments

**Figure 3.**
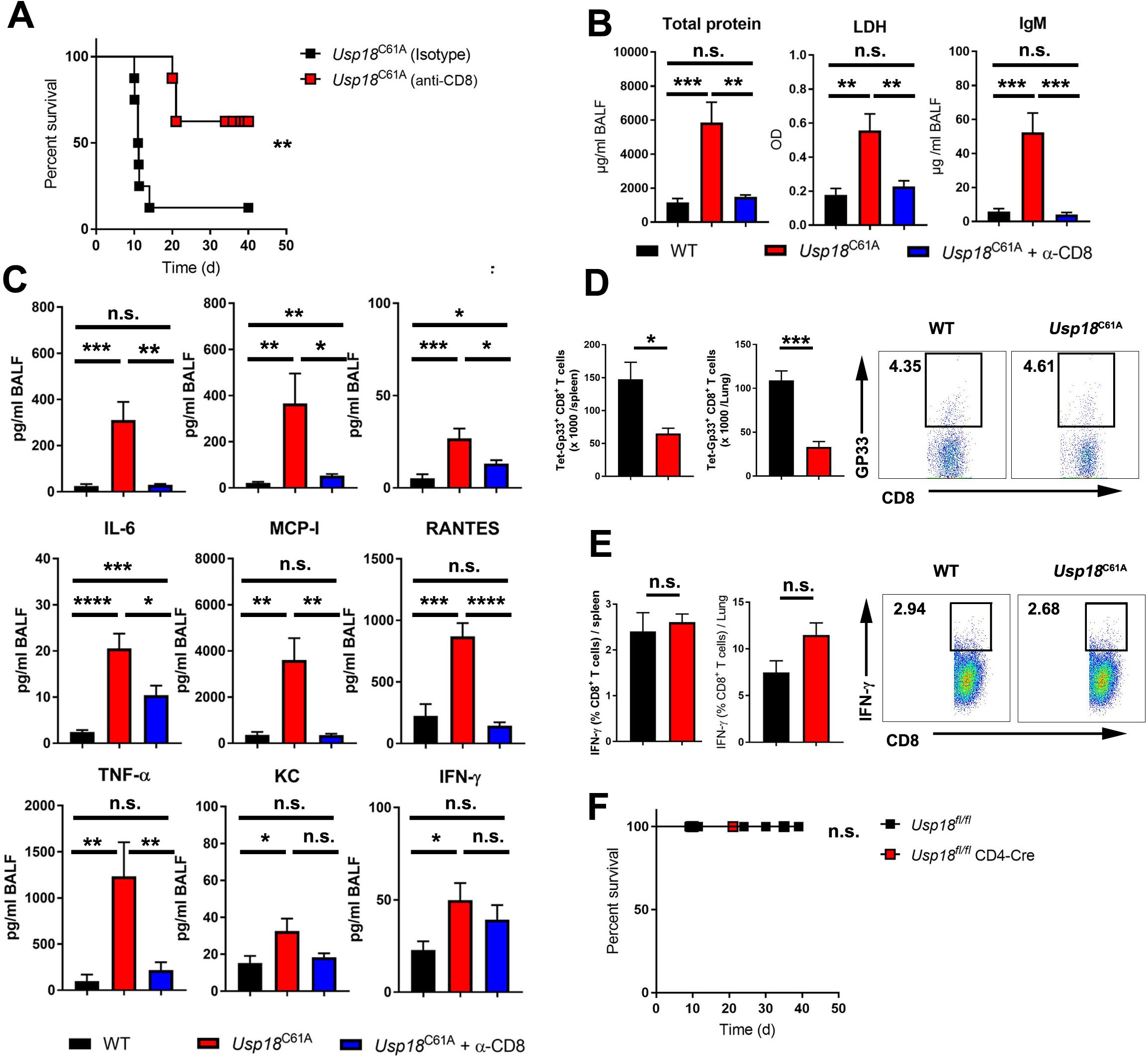
T cells mediates immunopathology in *Usp18*^C61A^ mice. (A) *Usp18*^C61A^ mice were treated with anti-CD8 or isotype antibodies at day -1 and 3. Mice were infected with 2 ×10^6^ PFU LCMV-Cl13 and survival was monitored (n = 6-8 mice/group). (B-C) *Usp18*^C61A^ mice were treated with anti-CD8 or isotype antibodies at day -1 and 3. Mice were infected, in addition to WT control, with 2 ×10^6^ PFU LCMV-Cl13 and 8 days post infection, (B) total protein, LDH and IgM were measured by BCA or ELISA assays (n = 8-10 mice/group). (C) indicated cytokines were measured in the BALF fluid (n = 8-10 mice/group). (D-E) WT and *Usp18*^C61A^ mice were infected with 2 x10^6^ PFU LCMV-Cl13. On day 8, (D) Total numbers and frequencies of LCMV-GP_33-41_ specific CD8 T cells were quantified by flow cytometry. (E) CD8^+^ T cells stimulated with GP_33-41_ peptide were intracellular stained for IFN-γ in the spleen and lung (n = 5 mice/group). (F) *Usp18*^*fl/fl*^ and *Usp18*^*fl/fl*^ x CD4-Cre were infected with 2 x10^6^ PFU LCMV-Cl13. Mice survival was monitored (n = 5-10 mice/group). n.s., not significant; *P < 0.05; **P < 0.01; ***P < 0.001 and ****P < 0.0001. Data are representative of at least two independent experiments

To determine if hyper-ISGylation was the principle driver of the above phenomena, we generated *Usp18*^C61A^ x *Uba7*^*-/-*^ mice, which are unable to either ISGylate or deISGylate, and assessed how these animals fared compared to *Usp18*^C61A^ or WT mice. The absence of *Uba7* in *Usp18*^*C61A*^ mice rescued production of most cytokines and chemokines to WT levels compared to *Usp18*^*C61A*^ mice following Cl13 infection, although Eotaxin and G-CSF levels were similar or elevated in *Usp18*^*C61A*^ x *Uba7*^*-/-*^ mice, respectively (**Figure 1G**). Despite restoration of many cytokine and chemokine levels in *Usp18*^C61A^ x *Uba7*^*-/-*^ compared to *Usp18*^*C61A*^ mice, simultaneous deletion of *Uba7* and a mutation in *Usp18* (C61A) did not result in dramatic reductions of markers signifying lung vascular leakage (**Figure 1H**), and, surprisingly, *Usp18*^*C61A*^ x *Uba7*^*-/-*^ mice succumbed to infection at rates similar to *Usp18*^*C61A*^ mice (**Figure 1I**). Together, these results indicate that hyper-ISGylation is not the underlying cause for the severe lung pathology observed in *Usp18*^*C61A*^ mice, even though certain immunologic manifestations are rescued in the absence of any ISGylation.

### Lack of the USP18 isopeptidase function leads to lung immunopathology and mortality in an ISG15-dependent manner

In addition to protein conjugation, ISG15 can also be secreted from cells and act as a cytokine-like molecule (D’Cunha et al., 1996; Knight and Cordova, 1991; Perng and Lenschow, 2018; Recht et al., 1991; Swaim et al., 2017). Thus, we asked if ablation of the enzymatic activity of USP18 would alter the levels of extracellular ISG15 in animals following Cl13 infection and if the extracellular pool of ISG15 could be responsible for the pathology. We first measured extracellular levels of ISG15 in the BALF following Cl13 infection. Interestingly, *Usp18*^C61A^ mice had significantly elevated levels of ISG15 in the BALF following Cl13 infection compared to WT mice (**Figure 2A**). Moreover, *Usp18*^*C61A*^ *x Uba7*^*-/-*^ mice had similar levels of ISG15 in BALF following Cl13 infection as *Usp18*^*C61A*^ mice (**Figure 2A**). Collectively, these results suggest that secreted ISG15 may be responsible for the severe pathology observed in Cl13-infected *Usp18*^*C61A*^ and *Usp18*^*C61A*^ *x Uba7*^*-/-*^ mice. Although IFN-I signaling is one of the strongest inducers of ISG15, additional stimuli such as lipopolysaccharide (LPS), retinoic acid and chemical toxins can induce ISG15 through interferon-dependent mechanisms (Kim et al., 2005; Liu et al., 2004; Malakhova et al., 2002; Perng and Lenschow, 2018). We next asked whether IFN-I signaling was required to produce the pathological manifestations observed in *Usp18*^C61A^ mice following Cl13 infection. Treatment of *Usp18*^C61A^ mice with anti-IFNAR1 antibody prior to Cl13 infection completely rescued survival (**Figure 2B**). To further confirm that ISG15 was necessary for the observed pathology, we infected *Usp18*^C61A^ x *Isg15*^*-/-*^ mice with Cl13. *Usp18*^C61A^ x *Isg15*^*-/-*^ showed only mild signs of morbidity, and 90% of *Usp18*^C61A^ x *Isg15*^*-/-*^ mice survived Cl13 infection (**Figure 2C**). Moreover, lack of ISG15 in *Usp18*^C61A^ mice alleviated pulmonary edema and alveolar wall thickening (**Figure 2D**). The rescue of lung pathology in *Usp18*^C61A^ x *Isg15*^*-/-*^ mice was accompanied by reduced vascular leakage into the airways indicated by lower levels of total protein, LDH and IgM in the BALF (**Figure 2E**). Furthermore, cytokine amplification observed in *Usp18*^C61A^ mice was largely reversed in *Usp18*^C61A^ x *Isg15*^*-/-*^ mice (**Figure 2F**). Together, our data demonstrate that ISG15 is required for the enhanced morbidity, mortality and pulmonary inflammatory pathology associated with enzymatically inactive USP18. This indicates that extracellular release of ISG15 potentiates the severe lung pathology and mortality observed.

### CD8 T cells are necessary for pulmonary immunopathology in *Usp18*^C61A^ mice

Infection of certain mouse strains like New Zealand Black (NZB), FVB/N and DBA-1 with Cl13 induces a robust T cell response and an acute CD8 T cell-mediated immunopathology causing lethality by day 9 post-infection (Baccala et al., 2014; Oldstone et al., 2018). The time at which *Usp18*^C61A^ mice succumbed to infection and the onset of cytokine amplification and lung vascular leakage closely correlated with the peak of the T cell-dominated adaptive immune response. Thus, we hypothesized that CD8 T cells may contribute to the severe inflammatory phenotype and the associated mortality in *Usp18*^C61A^ mice. To investigate this, we depleted *Usp18*^C61A^ mice of CD8 T cells prior to infection and assessed survival and lung inflammatory responses. CD8 T cell depletion in *Usp18*^C61A^ mice resulted in reduced mortality. Only ∼40% of CD8 T cell-depleted *Usp18*^C61A^ mice succumbed to the infection as compared to 90% of isotype control treated *Usp18*^C61A^ animals (**Figure 3A**). Moreover, depletion of CD8 T cells in *Usp18*^C61A^ mice significantly reduced lung leakage and cytokine storm following Cl13 infection compared to WT controls (**Figure 3B, C**). Interestingly, virus-specific T cell numbers were lower in the spleen and lung of *Usp18*^C61A^ mice than in WT mice (**Figure 3D**), and, upon re-stimulation, T cells of *Usp18*^C61A^ mice produced similar levels of intracellular IFN-γ (**Figure 3E**). These data suggest that, although CD8 T cells play a major role in driving the mortality observed in *Usp18*^C61A^ mice post-Cl13 infection, cytokine amplification, lung leakage and mortality observed in *Usp18*^C61A^ mice following Cl13 infection is not due to increases in virus-specific T cell numbers or functional capacity.

T cells are a major source of IFN-γ during Cl13 infection (Shaabani et al., 2016). We hypothesized that T cell-produced IFN-γ may promote the immunopathology observed in *Usp18*^C61A^ knock-in mice post-Cl13 infection. To test this, we treated Cl13 infected *Usp18*^C61A^ mice with an IFN-γ neutralizing antibody. Indeed, IFN-γ neutralization abolished lung leakage, cytokine storm and fatal outcome in *Usp18*^C61A^ mice (**Supplementary Figure 4A-C**). Further, both depletion of CD8 T cells and IFN-γ neutralization caused only slight but not significant reduction in the levels of free ISG15 in BALF of *Usp18*^C61A^ mice (**Supplemental Figure 4D**).

**Figure 4.**
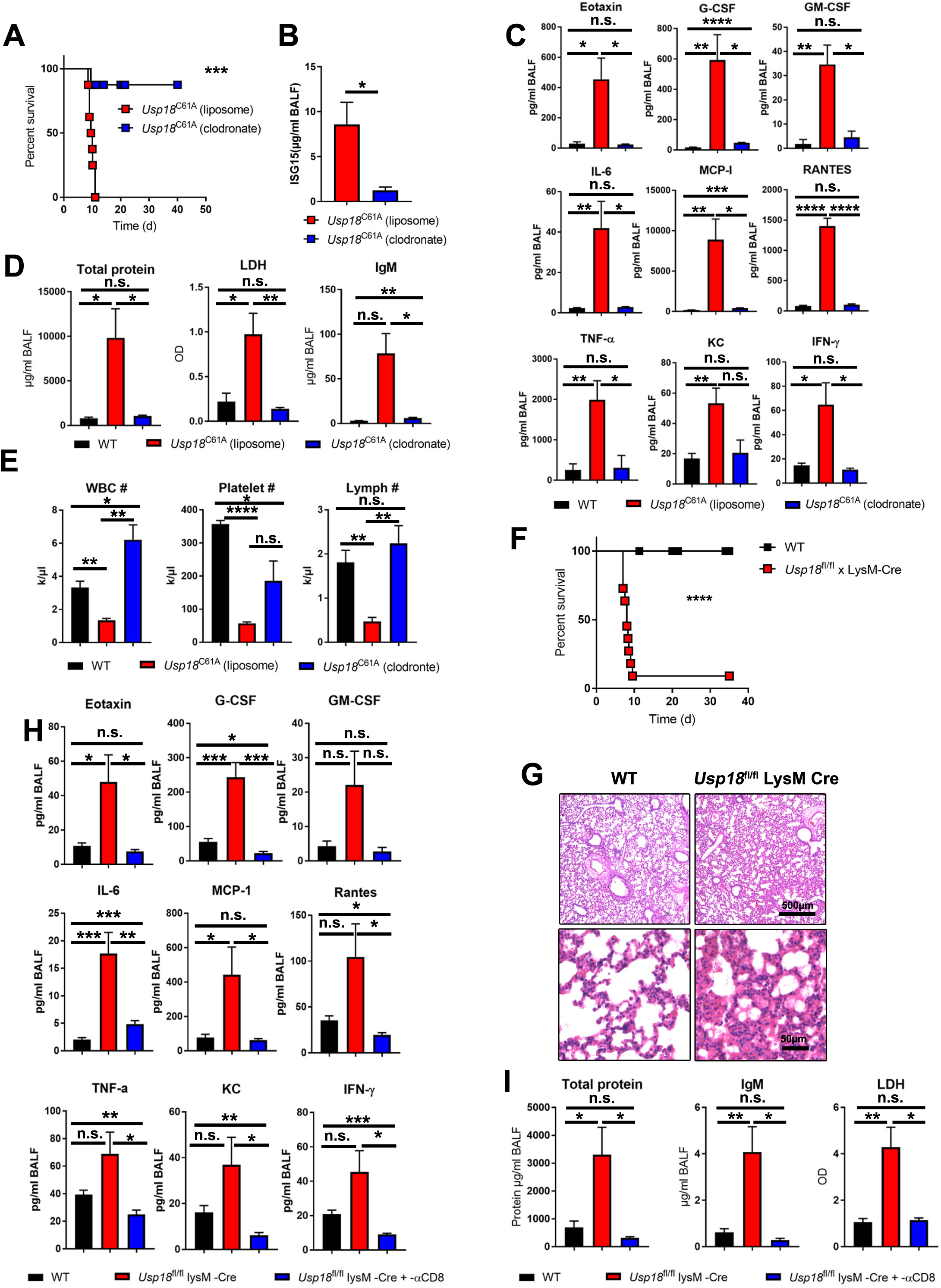
*Usp18* deletion in LysM^+^ cells reproduces the inflammatory pathology, cytokine amplification and mortality observed in *Usp18*^C61A^ mice. (A) *Usp18*^C61A^ mice were treated with empty or clodronate-loaded liposomes on day -1 and 4 pre infection with 2 ×10^6^ PFU LCMV-Cl13. Survival was monitored (n = 8 mice/group). (B-D) *Usp18*^C61A^ mice were treated with liposome or clodronate-liposome one day pre infection with 2 x10^6^ PFU LCMV-Cl13. On day 8. (B) ISG15 levels were measured in BALF by ELISA (C) Indicated cytokines. (D) total protein, LDH and IgM were measured in the BALF by BCA or ELISA assays (n = 4-5 mice/group). (E) *Usp18*^C61A^ mice were treated with empty or clodronate-loaded liposomes one day pre infection with 2 × 10^6^ PFU LCMV-Cl13. On day 5, White blood cells (WBC), platelets and lymphocytes were measured in the blood (n = 4-5 mice/group). (F) *Usp18*^*fl/fl*^ x LysM-Cre and WT mice were infected with 2 ×10^6^ PFU LCMV-Cl13, survival was monitored (n=7 mice/group). (G) Lung sections of WT and *Usp18*^C61A^ mice infected with 2 ×10^6^ PFU LCMV-Cl13 for 7 days were stained with hematoxylin and eosin (n = 3 mice/group). (H-I) *Usp18*^*fl/fl*^ x LysM-Cre treated with anti-CD8 or isotype antibodies at day -1 and 3 and WT mice were infected with 2 ×10^6^ PFU LCMV-Cl13. On day 6, (H) indicated cytokines (I) total protein, LDH and IgM were measured in the BALF (n = 8-9 mice/group). n.s., not significant; *P < 0.05; **P < 0.01; ***P < 0.001 and ****P < 0.0001. Data are representative of at least two independent experiments

The ability to lessen the pathology in *Usp18*^C61A^ mice by depletion of CD8 T cells prompted us to ask whether T cell-intrinsic deISGylation was necessary to restrain immunopathology following Cl13 infection. To address this question, we crossed *Usp18*^*fl/fl*^ mice to CD4-Cre mice; where USP18 is primarily deleted in total αβ T cells. *Usp18*^*fl/fl*^ x CD4-Cre displayed no overt signs of morbidity, and all animals survived Cl13 infection (**Figure 3F**). In addition, *Usp18*^*fl/f*l^ x CD4-Cre animals displayed no signs of lung vascular leakage or cytokine amplification compared to *Usp18*^*fl/fl*^ littermate controls (**Supplementary Figure 5A and B**). Furthermore, lack of USP18 in T cells did not increase free ISG15 levels in the BALF (**Supplementary Figure 5C**). To confirm our finding, we crossed P14 and SMARTA transgenic mice, which harbor LCMV-specific CD8 or CD4 T cells, respectively, to *Usp18*^C61A^ mice to obtain LCMV-specific *Usp18*^*C61A*^ knock-in CD8 or CD4 T cells. We transferred a mixture of *Usp18*^C61A^ or WT SMARTA and P14 T cells into WT mice. Adoptive transfer of *Usp18*^*C61A*^ knock-in CD4 or CD8 T cells did not result in similar morbidity and mortality observed in Cl13 infected *Usp18*^C61A^ mice (**Supplementary Figure 5D**). Collectively, our results strongly suggest that the excessive immunopathology observed in *Usp18*^C61A^ mice requires T cells, specifically T cell-produced IFN-γ, but is not related to T cell-intrinsic deletion of *Usp18*.

**Figure 5.**
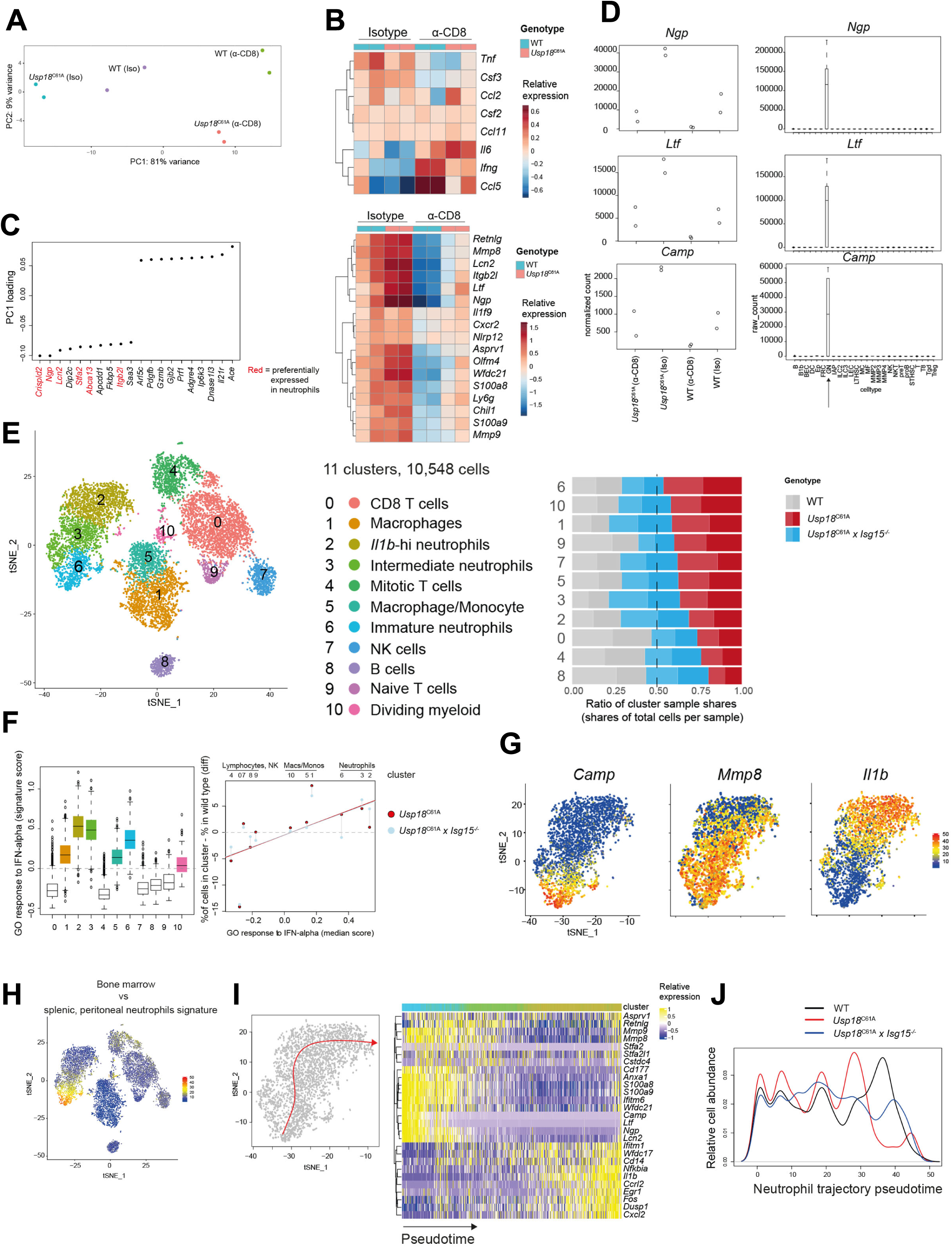
Transcriptome analysis of lung-infiltrating cells. (A-D) WT and *Usp18*^C61A^ mice were infected with Cl13 and treated with α-CD8 or isotype control. 7 days post infection, CD11b+ cells were purified from lung single cell suspension by flow cytometry, RNA isolated and sequenced using a ribo-depletion protocol. (A) Principal component analysis of total transcriptomes, top 2 components shown; (B) heatmap of relative expression of cytokine-encoding genes; (C) top 20 genes contributing to principal component 1 comprising 10 lowest and 10 highest loading value genes, neutrophil-enriched and neutrophil-specific genes marked in red; (D) left panels: expression of selected neutrophil markers per genotype, normalized counts shown; right panels: expression pattern of selected neutrophil markers in the Immgen dataset (GSE109125), cell type abbreviations as defined in GSE109125; GN, granulocyte neutrophils; (E-J) WT, *Usp18*^C61A^ and *Usp18*^C61A^ x *Isg15*^*-/-*^ mice were infected with Cl13, 7 days post infection, lungs collected infiltrating cells enriched for live singlets by flow cytometry, then subjected to single-cell transcriptome sequencing. (E) Dimensional reduction and clustering of cell transcriptomes (left panel); ratios of cells in cluster normalized by total cells per sample (right panel); (F) expression of the Gene Ontology gene set Response to interferon alpha per cluster (left panel); percentage change in cluster share of total cells in sample with respect to WT versus median expression of GO: response to interferon alpha gene set per cluster, mean values of biological replicates are shown, lines show linear regression for WT vs *Usp18*^C61A^, and WT vs *Usp18*^C61A^ x *Isg15*^*-/-*^; (G) heatmap of relative expression of neutrophil genes within the neutrophil clusters; (H) heatmap of expression of gene signature distinguishing bone marrow neutrophils from splenic and peritoneal neutrophils (derived from GSE109125); (I) differentiation trajectory within neutrophil clusters rooted at cluster 6, heatmap shows relative expression of neutrophil cluster marker genes along trajectory pseudotime; (J) density of cells along trajectory, biological replicates pooled by genotype, values normalized for total number of cells per genotype. Two biological replicates were used per genotype; further details see Materials and methods.

### Deletion of *Usp18* in LysM^+^ cells reproduces inflammatory pathology, cytokine amplification and mortality observed in *Usp18*^C61A^ mice

We next hypothesized that USP18 expression in innate immune cells may be essential for preventing disease following Cl13 infection. Given that T cells and IFN-γ were necessary for the pathology observed, we first assessed whether *Usp18* deletion in dendritic cells, the major cell population that presents antigen to T cells, was driving the immunopathology. We infected *Usp18*^*fl/fl*^ x CD11c-Cre mice, which mainly lack *Usp18* in CD11c^+^ dendritic cells, and found that *Usp18*^*fl/fl*^ x CD11c-Cre mice displayed no signs of overt pathology or mortality following Cl13 infection (**Supplementary Figure 6A**). We also observed no changes in lung vascular leakage or cytokine production in the BALF of *Usp18*^*fl/fl*^ x CD11c-Cre mice (**Supplementary Fig. 6B and C**), indicating that expression of *Usp18* in CD11c^+^ cells was not essential to curb pathology following Cl13 infection. Additionally, *Usp18*^*fl/fl*^ x CD11c-Cre mice did not show significantly elevated levels of extracellular ISG15 in the BALF compared to WT mice post-Cl13 infection (**Supplementary Figure 6D**). We speculated that other non-dendritic cell antigen presenting cells (APC) are activated by T cells in *Usp18*^C61A^ mice, leading to the hyper inflammatory state, cytokine storm and respiratory distress. To test this, we treated *Usp18*^C61A^ mice with clodronate-liposomes, which mainly deplete macrophages in addition to monocytes (Seiler et al., 1997; Sunderkotter et al., 2004). We found that clodronate treatment prevented mortality of *Usp18*^C61A^ mice (**Figure 4A**) which corresponded to a significant reduction in extracellular ISG15 levels in the BALF of *Usp18*^C61A^ mice after clodronate treatment (**Figure 4B**), strongly suggesting that these phagocytic cells are major sources of extracellular ISG15 following Cl13 infection. Additionally, clodronate-treated *Usp18*^C61A^ mice did not suffer from heightened cytokine amplification or lung vascular leakage (**Figure 4C, D**), and prevented cytopenia in *Usp18*^C61A^ mice (**Figure 4E**).

**Figure 6.**
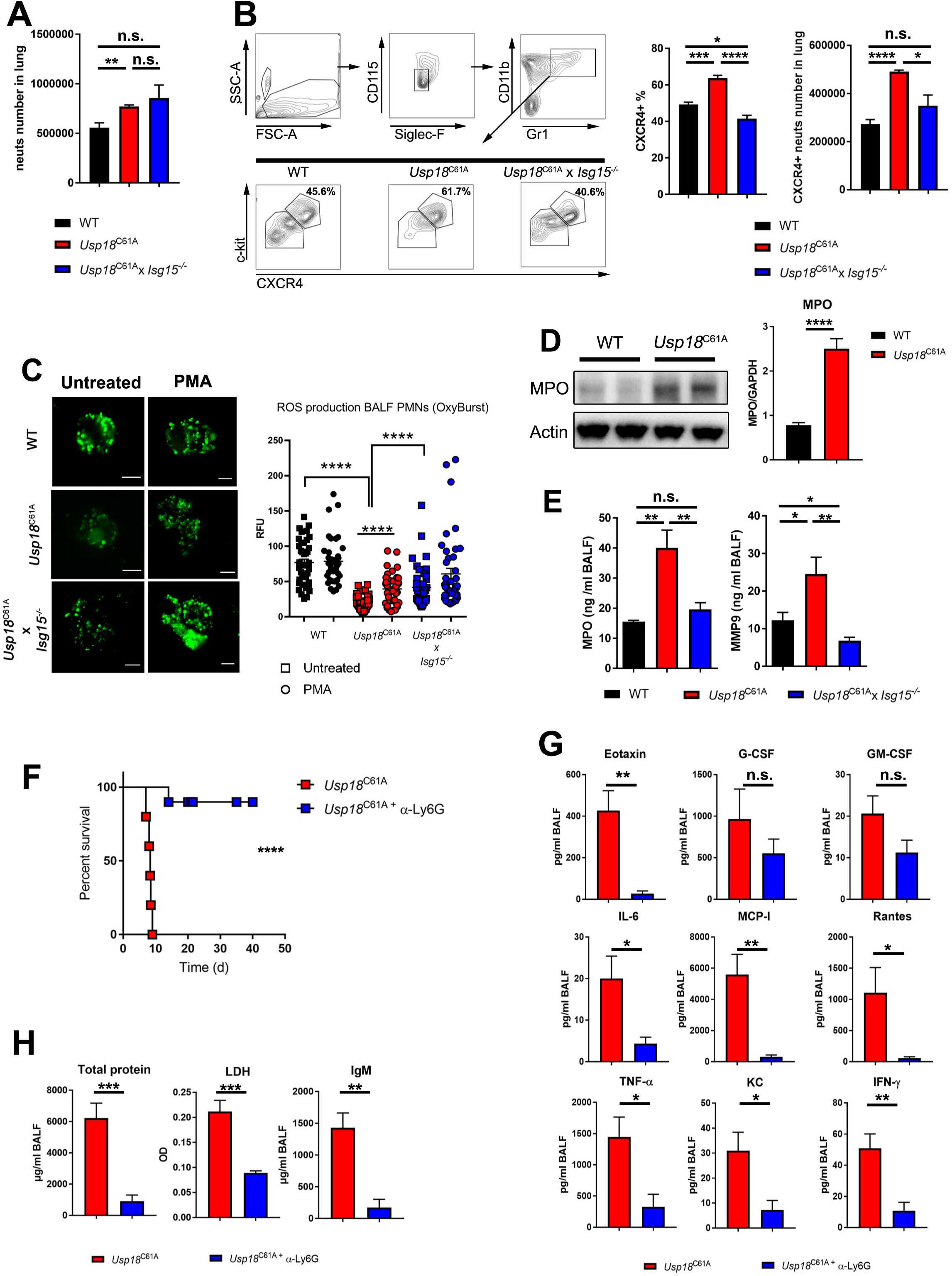
Lack of deISGylation during chronic infection increases migration of hyperactive neutrophils into lung leading to lethal immunopathology. (A-B) WT, *Usp18*^C61A^ and *Usp18*^C61A^ x *Isg15*^*–/–*^ mice were infected with 2 ×10^6^ PFU LCMV-Cl13. (A) Total numbers of CD11b^+^Ly6G^+^ neutrophils in the lung day 8 post-infection. (B) Flow cytometry staining of neutrophils in the lung on day 8 post-infection, frequencies and total numbers of c-kit^+^CXCR4^+^ neutrophils in the lung 8 days post-infection (n = 5 mice /group). (C) ROS production was measured in BALFs Polymorphonuclear cells with or without PMA stimulation (n = 37-46 cell/group). (D) WT and *Usp18*^C61A^ mice were infected with 2 ×10^6^ PFU LCMV-Cl13, myeloperoxidase (MPO) was assessed in the lung homogenates by western blot or (E) MPO and MMP9 were quantified by ELISA assay in BALF (n = 5 mice/group). (F) *Usp18*^C61A^ mice were infected with 2 ×10^6^ PFU LCMV-Cl13 and treated with anti-Ly6G or isotype antibodies at day 1 and 4 post-infection and monitored for survival (n = 5 Iso and 10 anti-Ly6G). (G-H) Mice were infected with 2 ×10^6^ PFU LCMV-Cl13 and 8 days post infection (G) indicated cytokines measured by multiplex assay and (H) total protein, LDH and IgM were measured in BALF by BCA and ELISA assay, respectively (n = 5 mice/group) n.s., not significant; *P < 0.05; **P < 0.01; ***P < 0.001 and ****P < 0.0001. Data are representative of at least two independent experiments

These findings led us to ask whether deletion of *Usp18* in myeloid cells, which include chlodronate targeted macrophages and monocytes, would result in pathological manifestations following Cl13 infection. To address this question, we infected *Usp18*^*fl/fl*^ and *Usp18*^*fl/fl*^ x LysM-Cre mice, which lack USP18 in monocytes, mature macrophages and granulocytes (Clausen et al., 1999), with Cl13. Following Cl13 infection, mice lacking *Usp18* in LysM^+^ cells exhibited significant mortality with 90% of *Usp18*^*fl/fl*^ x LysM-Cre^+^ animals succumbing to the infection by day 10 while no mortality was observed in littermate control *Usp18*^*fl/fl*^ animals (**Figure 4F**). Further, deletion of USP18 in LysM^+^ cells resulted in increased levels of extracellular ISG15 in BALF after Cl13 infection compared to Usp18^fl/fl^ litermates (**Supplementary Figure 7A**). Similar to Cl13 infection of *Usp18*^C61A^ mice, the increased mortality observed in *Usp18*^*fl/fl*^ x LysM-Cre^+^ mice was accompanied by extreme lung histopathology characterized by alveolar wall thickening, lung edema and increased mononuclear cell infiltrates (**Figure 4G**). We also observed increased vascular leakage and cytokine amplification in the BALF of *Usp18*^*fl/fl*^ x LysM-Cre compared to *Usp18*^*fl/fl*^ controls following Cl13 infection (**Figure 4H, I**). Moreover, *Usp18*^*fl/fl*^ x LysM-Cre suffered from hematological abnormalities with reduced white blood cells, lymphocytes and platelets (**Supplementary Figure 7B**). Overall, the pathological manifestations observed in *Usp18*^*fl/fl*^ x LysM-Cre mice mirrored those observed in *Usp18*^C61A^ mice (**Figure 1**). Interestingly, similar to what was observed in *Usp18*^C61A^ mice, depletion of CD8 T cells in *Usp18*^*fl/fl*^ x LysM-Cre mice prior to Cl13 infection diminished lung leakage and cytokine amplification in the BALF following Cl13 infection (**Figure 4 H-I**). These results show that mice lacking *Usp18* in myeloid cells phenocopy *Usp18*^C61A^ mice post-Cl13 infection, suggesting that lack of deISGylation in LysM^+^ cells results in CD8 T cell-mediated immunopathology during Cl13 infection.

### RNA-sequencing uncovers dysregulated neutrophil infiltration and differentiation

Our observation of increased BALF cytokines in *Usp18*^*C61A*^ mice associated with mortality prompted the question of which cell type(s) was responsible for the increased cytokine production. Increased cytokine levels in the lung could result from increases in cytokine production and secretion and/or higher numbers of cytokine-producing cells. To investigate the former, WT and *Usp18*^*C61A*^ mice were treated with CD8 T cell-depleting antibody or isotype control, and CD11b^+^ lung-infiltrating cells at the peak of morbidity were isolated and subjected to RNA-sequencing (**Supplementary Figure 8**). Principal component analysis reliably separated mice into experimental groups (**Figure 5A**). The bulk of variance could be accounted for by α-CD8 vs isotype treatment, but *Usp18*^*C61A*^ cells also separated from WT cells along the same component that separated α-CD8 from isotype-treated mice (**Figure 5A**). Interestingly, transcripts of cytokines with increased protein levels in the BALF were not upregulated, but some cytokines were downregulated in *Usp18*^*C61A*^ vs WT mice (**Figure 5B**). Moreover, cytokine transcripts increased in cells from α-CD8-treated mice, although these mice survived Cl13 infection. These results argue against increased cytokine transcription as a mechanism for the observed protein cytokine release in the BALF and agree with a previous study demonstrating a correlation between ISGylation and an increase in cytokine secretion (Radoshevich et al., 2015).

Interestingly, among the top-variance genes in this bulk RNA-seq dataset were many cell type-specific genes. Most notably, neutrophil-specific genes were upregulated in *Usp18*^*C61A*^ cells compared to WT (**Figure 5C**). The specificity of these genes to neutrophils was confirmed using a large consortium-generated dataset of mouse immune cell types (**Figure 5D**). Although CD8 T cell depletion dramatically reduced neutrophil transcript levels in WT cells, *Usp18*^*C61A*^ cells from α-CD8-treated mice exhibited levels similar to WT mice treated with an isotype control (**Figure 5D**). These results correlate with survival outcome in these groups, as *Usp18*^*C61A*^ isotype treated mice were more succumb to Cl13 infection (**Figure 3A**).

To gain a deeper understanding of the cell type dynamics associated with the observed lung pathology in infected *Usp18*^*C61A*^ knock-in mice, we performed single-cell RNA-seq of lung-infiltrating cells in WT, *Usp18*^*C61A*^ and *Usp18*^*C61A*^ *x Isg15*^*-/-*^ mice at the same infection timepoint (7 d.p.i.). Dimensionality reduction followed by clustering confidently identified major immune cell types from 3’ transcriptome data of 10,548 cells that passed quality control (**Figure 5E**). The major cell types by cell count were myeloid (neutrophils, macrophages, monocytes), with lymphocytes and NK cells accounting for a smaller fraction (**Figure 5E**). Comparing cluster composition by genotype revealed significant differences: *Usp18*^*C61A*^ mice showed a dramatic increase in neutrophil subsets compared to WT and *Usp18*^*C61A*^ x *Isg15*^-/-^ mice (**Figure 5E**). Further, *Usp18*^*C61A*^ mice also exhibited increased frequencies of other myeloid subsets and naïve T cells, whereas most B cells and other T cell subsets were underrepresented compared to WT mice.

Strikingly, there was a close correlation between relative levels of ISG expression and the relative increase in cell type frequency in *Usp18*^*C61A*^ vs. WT mice (**Figure 5F**). This suggests that cell types most sensitive to the C61A mutation are those experiencing a strong induction of *Usp18* and *Isg15* associated with ISG transcription. Although deletion of *Isg15* from *Usp18*^*C61A*^ mice led to distinct changes in infiltrating cell type frequencies with respect to WT controls (**Figure 5E-F**), the correlation between ISG expression and changes in percent infiltration remained (**Figure 5F**). Given that *Usp18* is strongly induced by interferon, this suggests that the catalytic region of USP18 may have additional, ISG15-independent functions.

The neutrophil subset specifically enriched in *Usp18*^*C61A*^ mice expressed elevated *Camp, Ltf* and *Ngp* transcripts, echoing the bulk RNA-seq results, and was low in *Il1b* and *Cxcl2* (**Figure 5G**). Comparison with published datasets demonstrated these *Ngp*-high neutrophils bear a transcriptional signature of secondary granule genes associated with bone marrow-derived, immature neutrophils (**Figure 5H**). mRNA expression of these genes ceases in peripheral mature neutrophils (Evrard et al., 2018). Trajectory analysis revealed a continuous transcriptome transition between these neutrophil subsets, suggesting immature neutrophils can differentiate into more mature subtypes *in situ* (**Figure 5I**). Overall, there was a major shift in the neutrophil population towards immature neutrophils in *Usp18*^*C61A*^ mice whereas *Usp18*^*C61A*^ *x Isg15*^-/-^ mice accumulated more differentiated, *Mmp8*^+^ *Il1b*-intermediate neutrophils compared to WT mice (**Figure 5E, J**). The dichotomous effect of the *Usp18*^*C61A*^ and *Isg15*^*-/-*^ alleles on neutrophil subtypes suggests ISG15 secretion could regulate neutrophil activation and/or differentiation in addition to migration.

### Lack of deISGylation increases CXCR4^+^ neutrophils in lung leading to lethal immunopathology

Given our RNA sequencing results, we expected to observe changes in neutrophil subsets in the lungs in *Usp18*^C61A^ knock-in mice following Cl13 infection. We initially measured total numbers of neutrophils in lungs of Cl13-infected WT and *Usp18*^C61A^ mice and observed a modest but significant increase in total lung neutrophils in *Usp18*^C61A^ mice compared to WT controls (**Figure 6A**). More comprehensive flow cytometry staining revealed significant increases in the frequencies and total numbers of CXCR4^+^ neutrophils in the lung of *Usp18*^*C61A*^ mice compared to WT controls (**Figure 6B**) and agreed with the gene expression of *Cxcr4* in CD11b^+^ lung-infiltrating cells (**Supplementary Figure 9**). Interestingly, total neutrophil numbers were also increased in *Usp18*^*C61A*^ x *Isg15*^*-/-*^ mice compared to WT controls following Cl13 infection (**Figure 6A**). However, deletion of *Isg15* from *Usp18*^*C61A*^ mice restored the frequencies and total numbers of CXCR4^+^ neutrophils in the lung to levels similar to or lower than those observed in WT mice following Cl13 infection (**Figure 6B**). Analysis of neutrophil reactive oxygen species (ROS) production by BALF neutrophils from WT, *Usp18*^*C61A*^ and *Usp18*^*C61A*^ x *Isg15*^*-/-*^ revealed reduced ROS production *ex vivo* from unstimulated and stimulated *Usp18*^*C61A*^ neutrophils compared to WT neutrophils (**Figure 6C**), suggesting at the time of analysis *Usp18*^*C61A*^ neutrophils appear exhausted. Notably, *Usp18*^*C61A*^ x *Isg15*^*-/-*^ neutrophils produced WT-levels of ROS both before and after PMA stimulation (**Figure 6C**). Neutrophils release various granule proteins including myeloperoxidase (MPO), from azurophilic granules, which can damage lung-resident epithelial and endothelial cells (Haegens et al., 2008; Schurmann et al., 2017). We measured MPO in total lung homogenates by western blot and found elevated levels in *Usp18*^C61A^ compared to WT lungs. (**Figure 6D**). Additionally, MPO and MMP9 (gelatinase granules) levels were higher in the *Usp18*^C61A^ BALF compared to WT BALF (**Figure 6E**), suggesting that MPO and MMP9 are secreted into the lung milieu by infiltrating neutrophils. Importantly, the levels of MPO and MMP9 in the BALF of *Usp18*^*C61A*^ x *Isg15*^*-/-*^ mice were restored to similar levels as WT controls (**Figure 6E**). Moreover, depletion of either CD8 T cells or IFN-γ restored the levels of MPO and MMP9 in *Usp18*^C61A^ mice similar to those observed in isotype treated WT mice (**Supplementary Figure 10**), indicating that T cell-produced IFN-γ promotes MPO and MMP9 production or secretion by lung neutrophils in *Usp18*^*C61A*^ mice following Cl13 infection. To show that neutrophils are the primary mediators of immunopathology in the lung of *Usp18*^C61A^ mice during Cl13 infection, we depleted neutrophils using an anti-Ly6G antibody. Neutrophil depletion rescued survival in Cl13-infected *Usp18*^C61A^ mice, with 90% of neutrophil depleted *Usp18*^*C61A*^ mice surviving compared to 10% survival observed in isotype control-treated *Usp18*^*C61A*^ mice (**Figure 6F**). Moreover, neutrophil depletion significantly reduced lung leakage and cytokine storm in *Usp18*^C61A^ mice (**Figure 6G, H**). Together, we conclude that lack of deISGylation in LysM^+^ cells during persistent viral infection ultimately results in neutrophil hyperactivation and excessive release of azurophilic and gelatinase granule cargoes, including MPO and MMP9, which contributes to lung damage and death. Furthermore, our data suggests that high levels of extracellular ISG15 produced by LysM^+^ cells promotes the accumulation of CXCR4^+^ neutrophils in the lung following Cl13 infection and the resultant immunopathology.

## Discussion

IFN-I signaling induces the expression of hundreds of genes, with *Isg15* amongst the most dynamically upregulated (Perng and Lenschow, 2018). Despite the “ubiquitous” conjugation of ISG15 to many host proteins, the biological consequences of proper regulation of ISG15 conjugation and deconjugation in immune cells has been understudied. The ability of cells to produce extracellular ISG15 has been known for many years (D’Cunha et al., 1996; Knight and Cordova, 1991; Perng and Lenschow, 2018; Recht et al., 1991; Swaim et al., 2017). However, the mechanism of ISG15 secretion into the extracellular environment is currently unknown. Our study is the first to report a role for USP18 isopeptidase function in regulating extracellular levels of ISG15 *in vivo*. This finding is further supported by a previous study where elevated levels of free ISG15 in cells and tissues of *Usp18*^*C61A*^ mice were reported (Ketscher et al., 2015). Specifically, our results demonstrate that proper ISGylation/deISGylation in LysM^+^ cells regulates extracellular ISG15 levels, and perturbation of this pathway results in lung immune pathology, cytokine amplification and mortality following persistent Cl13 infection. Thus, we believe that LysM^+^ cells are major sources of extracellular ISG15 following persistent LCMV infection. The exact mechanism by which the inability to deISGylate results in higher levels of extracellular ISG15 is uncertain but we can offer some hypotheses: one possibility is that USP18 binding to ISG15 in cells regulates levels of free ISG15, possibly through targeted degradation. A second hypothesis is that cellular ISGylated substrates become saturated in *Usp18*^C61A^ knock-in cells and the lack of potential ISGylated protein substrates results in increased quantities of free ISG15 which can be secreted. The third possibility is that strong ISGylation leads to increased cell death, resulting in the release of free ISG15. Further cellular and biochemical studies will be required to understand how the enzymatic function of USP18 regulates extracellular ISG15.

Elevated levels of IFN-I have been linked to severe lung inflammation for a number of viruses that replicate in the respiratory tract including Orthomyxoviruses, Coronaviruses and Arenaviruses. Importantly, genetic deletion, pharmacological inhibition or antibody neutralization of IFN-I signaling can alleviate severe pulmonary pathologies in animal models of Influenza, SARS-CoV and LCMV infections (Baccala et al., 2014; Channappanavar et al., 2016; Davidson et al., 2014; Davidson et al., 2015; Teijaro, 2016; Teijaro et al., 2011). Given that *Isg15* is a highly induced ISG, it is possible that inhibition of IFN-I signaling results in reduced ISG15 production/secretion, resulting in reduced pathological outcomes during these infections. More evidence for a causal role of ISG15 in promoting inflammatory pulmonary pathology can be inferred from humans with mutations causing loss of function of *USP18* or *ISG15*. In humans, USP18 loss-of-function mutations result in a pathological condition called Pseudo-TORCH syndrome. These patients suffer from severe acute respiratory distress, cytopenia, hemorrhage, and septic shock (Dauphinee et al., 2014; Lindner et al., 2007) with the majority succumbing to the condition in late gestation or early infancy. Despite the increased levels of ISGylation observed in these patients, their disease was primarily attributed to elevated IFN-I signaling and the resultant interferonopathy (Alsohime et al., 2020; Meuwissen et al., 2016). Data presented herein suggests that dysregulated ISGylation/deISGylation in the absence of USP18 can be causal in aberrant immune inflammation in the lung. This hypothesis is further supported in humans, where ISG15 has been reported to stabilize USP18 and *ISG15*-null patients exhibit strong IFN-I signaling but display much less severe symptoms than *USP18*-loss of function patients, suggesting that secreted ISG15 may significantly contribute to symptoms observed in *USP18*-null patients. Furthermore, if free ISG15 contributes to disease pathogenesis in *USP18*-null patients, ISG15 levels can have prognostic value in assessing the potential for severe pathological outcomes in *Usp18*-deficient patients or following viral infections. Moreover, antibody neutralization or pharmacological inhibition of ISG15 signaling may be a viable means to alleviate pathology and mortality in *USP18*-deficient patients or during viral infection where IFN-I signaling correlates with severe pathological outcomes.

Previously it was demonstrated that extracellular ISG15 promotes neutrophil chemotaxis *in vitro* (Owhashi et al., 2003). However, whether ISG15 acts as a chemoattractant *in vivo* remains unclear. Additionally, it was reported that ISG15 is present in neutrophil granules and can be secreted following bacterial exposure (Bogunovic et al., 2012). Furthermore, patients that lack ISG15 produce less IFN-γ and are unable to control mycobacterial infections (Bogunovic et al., 2012), highlighting the importance of ISG15 in controlling bacterial infection in humans. Our study is the first to report a role for the enzymatic function of USP18 in regulating extracellular levels of ISG15. Specifically, we demonstrate that inability to remove ISG15 from cellular protein substrates promotes elevated levels of extracellular ISG15, resulting in the accumulation of CXCR4^+^ neutrophils in the lungs following persistent virus infection. Moreover, the neutrophil infiltrate in *Usp18*^*C61A*^ mice was highly pathogenic, producing elevated levels of MPO and MMP9, promoting cytokine storm and death. Importantly, complete deletion of *Isg15* from *Usp18*^C61A^ mice reversed the accumulation of CXCR4^+^ neutrophils and the associated pathological outcomes. CXCR4 expression on neutrophils promotes retention and survival of neutrophils in inflamed tissues (Isles et al., 2019). Elevated levels of neutrophil CXCR4 expression and its ligand CXCL12 has been found in patients with severe lung inflammation in COPD and cystic fibrosis patients (Carevic et al., 2015; Hartl et al., 2008), and CXCR4^+^ neutrophils are elevated in late stage LPS-induced lung injury (Yamada et al., 2011). Thus, CXCR4^+^ neutrophils are found in scenarios of severe lung pathology. Moreover, the fact that neutrophil depletion rescued survival in *Usp18*^C61A^ mice further demonstrates that the neutrophil population recruited to the lung is causal in promoting the inflammatory pathology, morbidity and mortality observed in *Usp18*^*C61A*^ mice following persistent virus infection. The role of neutrophils in mediating pathology during Arenavirus infection has not been studied in detail, and our data suggest that neutrophils may play a causal role in disease warranting further investigations in humans and animal models of Arenavirus infection.

The requirement of CD8 T cells for inflammatory pathology, morbidity and mortality in *Usp18*^C61A^ mice suggests that T cells and neutrophils engage in significant cross talk which modulates neutrophil maturation or inflammatory states. We still do not understand exactly how T cells and IFN-γ are affecting neutrophil differentiation/migration/activation. Extracellular ISG15 has been reported to signal on NK cells and lymphocytes to induce the production of IFN-γ, therefore it is possible that ISG15 and IFN-γ act in a feed forward loop to promote amplification of pathological neutrophils during viral infection. Previously, it was reported that CD8 T cells promoted a massive secondary recruitment of pathogenic monocytes and neutrophils into the meninges following LCMV infection which resulted in seizures and rapid lethality, although the mechanism remains incompletely defined (Kim et al., 2009). We report here that IFN-γ is a key player in death of *Usp18*^C61A^ mice following Cl13 infection, as IFN-γ neutralization was superior at reducing inflammation, morbidity and mortality compared to CD8 T cell depletion. The production and release of IFN-γ by T cells can prime neutrophil functions such as phagocytosis, cell killing, cytokine release and nitric oxide production (Ellis and Beaman, 2004). Most likely, IFN-γ induced by T cells leads to hyper-activation of *Usp18*^C61A^ neutrophils. In support of this hypothesis, neutralization of IFN-γ reduced BALF levels of MPO and MMP9 in *Usp18*^C61A^ mice. However, the extent to which neutrophil-T cell crosstalk promotes the inflammatory pathology observed in *Usp18*^C61A^ mice following Cl13 infection remains incomplete and is an exciting area for future study. Taken together, our study is the first to implicate a pathological role of extracellular ISG15 during viral infection and may have broad implications for diagnosis and treatment of severe respiratory pathology in interferonopathies, viral infections and additional IFN-I-driven diseases.

## Materials and Methods

### Mice, Virus

C57BL6/J, *Usp18*^*–/–*^, *Usp18*^fl/fl^ CD11c-Cre, *Usp18*^fl/fl^ LysM-Cre, *Usp18*^fl/fl^ CD4-Cre, *Usp18* C61A/C61A, *Usp18* ^C61A/C61A^ x *Isg15*^*–/–*^, *Usp18* ^C61A/C61A^ x *Uba7*^*–/–*^, P14 and SMARTA male and female mice (7-9 weeks of age) were used. *Usp18*^fl/fl^ mice were provided by Prof. Marco Prinz, *Usp18* C61A/C61A mice were provided by Dr. Klaus-Peter Knobeloch. *Usp18*^*–/–*^, *Uba7*^*–/–*^, and *Isg15*^*–/–*^ mice were provided by Prof. Dong-Er Zhang. *Usp18*^*–/–*^ were crossed with 129SVE-M129S6/SvEvTac purchased from Taconic laboratory for seven generations and mated heterozygous x heterozygous with WT littermates used as controls for all experiments. CD11c-Cre, CD4-Cre and LysM-Cre mice were purchased from Jackson laboratory. Except *Usp18*^*–/–*^ mice, all strains were maintained on C57BL6/J background. Mice were maintained in pathogen-free conditions and handling conforms to the requirements of the National Institutes of Health and the Scripps Research Institute Animal Research Committee. LCMV strain Armstrong or clone 13 were propagated on BHK cells and mice were injected with 2 x10^6^ PFU intravenously for LCMV-Cl13 or 2×10^5^ intraperitonially for LCMV-Arm. Mice were monitored daily for morbidity either they succumbed on their own or were sacrificed when they became excessively moribund, by losing >25% of weight accompanied with labored breathing.

### Antibodies and clodronate treatment

Mice were treated intraperitoneally with 1 mg of anti-IFNAR-1 clone MAR1-5A3 (Leinco; MO, USA), isotype IgG1 (Leinco) one day prior to LCMV infection. Mice were given 1 mg of anti-CD8 clone YTS 169.4 (Leinco) on day-1 and 3. For IFN-γ depletion, mice were treated with anti IFN-γ clone XMG1.2 (Leinco) 500µg i.p on day 1, 4 and 9 post infection. For macrophages and monocytes depletion, mice were treated intravenously with 200 µl of liposome-clodronate or liposome on day -1 and 4. For Ly6G depletion, mice were treated with 100µg of anti-Ly6G clone RB6-8C5 (Leinco) on day 1, 4 and 9 post infection.

### Total RNA extraction, c-DNA synthesis, and quantitative real-time PCR

RNA was isolated from splenocytes or sorted cells with TRIzol (Thermo Fisher) as described by the manufacture’s protocol. The RNA was reverse-transcribed into cDNA with the Quantitect Reverse Transcription Kit (Qiagen, Germany). Gene expression analysis was performed with assays from Qiagen: glyceraldehyde 3-phosphate dehydrogenase (*GAPDH*; QT01658692), *Irf7* (QT00245266), *Prkr* (QT00162715), *Usp18* (QT00167671), *and* Stat2 (QT00160216). Relative quantities (RQs) were determined with the equation RQ = 2^−ddCt^.

### Gel Electrophoresis and Immunoblotting

Cells were collected and lysed on ice for 30 min in 1X radioimmunoprecipitation assay (RIPA) buffer plus (Cell Signaling Technology) with protease and phosphatase inhibitor cocktail (Thermo Scientific). The lysate was centrifuged, and the supernatant was collected. Laemmli loading buffer (Biorad) with 0.05% BME was added to each sample. Samples were then heat denaturated for 5 min at 95 c° before loaded into the NuPAGE 4-12% (vol/vol) Bis-Tris SDS gels and 3-(N-morpholino) propanesulfonic acid buffer. The gels were run for 1 h at 180 V and were then transferred onto PVDF or nitrocellulose membranes using the Trans-Blot TURBO semidry transfer system from Bio-Rad.

The membrane was blocked for 1 h and then probed overnight with 1:1,000 of primary antibody in blocking solution at 4 °C overnight. The membrane was then washed three times for 10 min with Tris-buffered saline-tween (TBST). After the wash, the membrane was incubated with the corresponding Ab (HRP-conjugated) for 1 h at room temperature at a 1:5,000 dilution. The membrane was washed three times for 10 min in 1× TBST. Finally, the blots were overlaid with Clarity max ECL (Biorad) and imaged on the Biorad ChemiDoc.

### ELISAs

Multiplex ELISA (Biorad; CA, USA) was performed to detect cytokines in BALF. IFN-α and -β levels were measured in the serum according to PBL manufacturer’s instructions. Total proteins in BALF were measured using Pierce BCA protein assay kit (Thermo Scientific). IgM were measured by Mouse IgM quantification set (Bethyl laboratories). LDH was measured using cytotox96 Non-radioactive cytotoxicity assay (Promega).

ISG15 ELISA was performed by coating plate with diluted rabbit anti mouse ISG15 (PA5-17461; Thermo Scientific) 1:1000 in PBS overnight. Next day, plates were blocked with FBS 10 % in PBS for 2 hours. Afterward, samples were applied for 1 hour. Free ISG15 levels were detected by house made rat anti mouse ISG15 6µg /ml followed by anti-rat HRP and developed with TMB. Washing with PBS/Tween was performed between each step.

The concentration of gelatinase B (MMP-9) and myeloperoxidase (MPO) in the BALF was analyzed using mouse-specific MMP-9 or MPO ELISA duo kits (R&D), respectively.

### T cells transfer

CD4 or CD8 T cells were isolated from spleen using magnetic beads as described by manufacturer’s protocol (STEMCELL technologies). 4000 cells in ratio 1:1 were transferred into WT mice 1 day prior to LCMV-Cl13 infection.

### Histology

Histological sections were performed on formalin-fixed lungs. Hematoxylin and Eosin staining was performed in histology core in Scripps Research institute.

### FACS

Spleens or lungs were digested with collagenase from *Clostridium histolyticum* type IV (Sigma), deoxyribonuclease I from bovine pancreas grade II (Roche, IN, USA), and trypsin inhibitor type II-S soybean. Cells were washed with phosphate-buffered saline (Thermo Fisher Scientific) and stained with anti-CD11c (clone N418; BioLegend, CA, USA), anti-CD11b (clone M1/70; BioLegend), anti-CD8a (clone 53-6.7; BioLegend), anti-Ly6G (clone RB6-8C5; BioLegend), and anti-Ly6C (clone HK1.4; BioLegend), anti-CD19 (clone 6D5; BioLegend), anti-CD3 (clone 17A2; BioLegend). For measurement of intracellular cytokines, cells were stimulated with LCMV-GP33, fixed with 2% formaldehyde and permealized with BD Perm/Wash buffer (Fischer) and stained with anti IFN-γ (clone XMG1.2; BioLegend) and anti-GM-CSF (clone MP1-22E9: BioLegend).

### Bulk RNA-sequencing

RNA was isolated from flow cytometry purified cells using RNeasy Mini kit (Qiagen, Hilden, Germany) with on-column DNase digest. RNA quality was assessed using the Bioanalyzer Nano chip (Agilent Technologies, Santa Clara, CA) and sequencing libraries prepared using the NEB Next Ultra II Directional RNA kit (NEB, Ipswich, MA) then sequenced to a depth of at least 20 M single-end 75 bp reads using NextSeq 500 (Illumina, San Diego, CA).

### Bulk RNA-sequencing analysis

Demultiplexing was performed using bcl2fastq (version 2.17.1.14). Quality control of FASTQ files was performed using fastQC version 0.11.9 (Andrews, 2010). Reads were mapped to the mouse genome (GRCm38 assembly with Ensembl v95 transcriptome annotations) and gene counts calculated using STAR version 2.7.1a (Dobin et al., 2013) using the following parameters: --quantMode GeneCounts –outSAMtype None –sjdbGTfile /Ensembl/Mus_musculus.GRCm38.95.gtf –sjdbOverhang 74. Statistical normalization and differential expression analysis were performed using R programming language version 3.6.1 (Team, 2008) and the R package DESeq2 version 1.26.0 (Love et al., 2014). Reference data for panel 6D were obtained from re-aligned reads from GSE109125 (Yoshida et al., 2019).

### Single-cell RNA-sequencing

Single live cells from individual mice were isolated by flow cytometry and labeled with TotalSeq-B cell hashing antibodies (Biolegend, San Diego, CA). Labeled cells were subjected to 3’ version 3.1 transcriptome library preparation using the Chromium 10X system (10X Genomics, Pleasanton, CA) following manufacturer’s instructions. Libraries were pooled and sequenced using the recommended 10X protocol for Illumina on the Mid Output flow cell for NextSeq 500 (Illumina).

### Single-cell RNA-sequencing analysis

Sequencing reads were demultiplexed using cellranger mkfastq version 3.1.0 (10X Genomics). Following quality control with fastQC version 0.11.9, cell identification, read extraction, alignment, barcode read identification and read counting were performed using cellranger count version 3.1.0 (10X Genomics). GRCm38 genome and Ensembl annotations version 95 were used as reference. Cell demultiplexing, normalization, clustering and dimensionality reduction were performed using the R package Seurat version 3.1.3 (Stuart et al., 2019). Basic plotting and statistical analyses were performed using R version 3.6.2 (Team, 2008). Trajectory analysis was performed using R package slingshot version 1.4.0 (Street et al., 2018). Heatmaps were plotted using R packages pheatmap version 1.0.12 (Kolde, 2013) and Complex Heatmap version 2.2.0 (Gu et al., 2016). For cluster ratios in Figure 6E, ratio of each sample was calculated using the as follows: sample cluster fraction = number of cells in cluster from sample / number of total cells in sample. Relative sample cluster fraction = sample cluster fraction / sum of sample cluster fractions for all samples. Gene Ontology signature for panel 6F was obtained from geneontology.org in March 2020.

### Immunofluorescence analysis

Neutrophils were seeded on untreated coverglass using a 4-Chamber 35 mm dish with 20 mm microwells (Cellvis LLC Cat # D35C4-20-1.5-N) and incubated at 37°C for 30 min, then fixed with 3.7% paraformaldehyde for 8 min, permeabilized with 0.01% saponin, and blocked with 1% BSA in PBS. The samples were labeled with the indicated primary antibodies overnight at 4°C in the presence of 0.01% saponin and 1% BSA. Samples were washed and subsequently incubated with appropriate secondary antibodies for 2 hours at room temperature. Neutrophil nuclei were stained with 4, 6-diamidino-2-phenylindole, dihydrochloride (DAPI) for 15 min at 21°C, washed with PBS and samples were subsequently mounted using Fluormount G. For the analysis of reactive oxygen species, BALF neutrophils were seeded on coverglass as above and ROS was analyzed using Oxy-burst (Carboxy-H2DCFDA). Samples were analyzed with a Zeiss LSM 880 laser scanning confocal microscope attached to a Zeiss Observer Z1 microscope at 21 □C, using a 63× oil Plan Apo, 1.4 numerical aperture (NA) objective. For the comparative quantification of ROS production between BALF neutrophils from the various colonies, gain and laser power were maintained invariant throughout the analysis.

### Statistics

Data are expressed as mean ± SEM Unpaired two-tailed student’s *t*-test was calculated using GraphPad Prism to perform statistically comparison between groups. Statistical comparison of several groups was performed by two-way analysis of variance (ANOVA). Log-rank (Mantel-Cox) test was used for survival study. The level of statistical significance was set at n.s. not significant, *\* P* < 0.05, *\*\* P* < 0.01, *\*\*\* P* < 0.001 or *\*\*\*\* P* < 0.0001.

## Acknowledgments

We want to thank Seema Saini, Nisha Rao and Jessica Van Leeuwen from the Department of Animal Resources and FACS Core in Scripps Research Institute for technical support.

## Author contributions

N.S and J.R.T designed and performed experiments, analyzed results and wrote the manuscript. N.S and J.Z performed statistical analysis. J.Z, N.N, Z.H, D.L, J.J and S.C performed experiments. N.H, K.A, M.P, K-P.K, S.C and D-E.Z discussed results.

## Supplemental Figures

**Supplementary Figure 1** (A-B) WT and *Usp18*^*–/–*^ mice were infected with 2×10^6^ PFU of LCMV-Cl13. 6 days post infection, (A) Indicated cytokines were quantified by multiplex assay and (B) total protein and LDH were quantified in the BALF by BCA and ELISA assays (n = 5-7 mice/group). n.s., not significant; *P < 0.05 and **P < 0.01.

**Supplementary Figure 2** (A-B) WT and *Usp18*^C61A^ mice were infected with 2 ×10^6^ PFU LCMV-Cl13. After 2 days, ISGs were quantified in the spleen (A) and lung (B) by RT-PCR (n = 4 mice/group). (C) WT and *Usp18*^C61A^ mice were infected with 2 ×10^6^ PFU LCMV-Cl13 and IFN-α and β were quantified by ELISA in the serum at indicated time points. (n = 4 mice/group).

**Supplementary Figure 3** WT and *Usp18*^C61A^ mice were infected with 2 x10^6^ PFU LCMV-Cl13. Viral titers were quantified in the indicated organs on day 5 and day 7 by focus forming assay (n = 4-5 mice/group).

**Supplementary Figure 4** (A) *Usp18*^C61A^ mice treated with 500ug of anti-IFN-γ or isotype antibodies at day 1, 4 and 8. Mice were infected with 2 x10^6^ PFU LCMV-Cl13 and survival was monitored (n = 5-6 mice/group). (B-C) *Usp18*^C61A^ mice treated with anti-IFN-γ or isotype antibodies at day 1 and 4. Mice were infected with 2 x10^6^ PFU LCMV-Cl13 and 8 days post infection and (B) indicated cytokines (n = 4-5).were quantified by multiplex assay or (C) total protein, IgM and LDH were quantified in BALF by BCA or ELISA assay (n = 4-5 mice/group). (D) WT, *Usp18*^C61A^ and *Usp18*^C61A^ mice treated with anti-IFN-γ or anti-CD8 were infected with 2 x10^6^ PFU LCMV-Cl13. 8days post infection, ISG15 levels were measured in the BALF (n = 4-5)

**Supplementary Figure 5** (A-C) *Usp18*^*fl/fl*^ and *Usp18*^*fl/fl*^ x CD4-Cre were infected with 2 x10^6^ PFU LCMV-Cl13 and 8 days post infection, (A) total protein, LDH and IgM were quantified by BCA and ELISA assay (B) indicated cytokines quantified my multiplex assay and (C) ISG15 levels were quantified in the BALF by ELISA (n = 4-5 mice/group). (D) 2000 P14 CD8 T cells x *Usp18*^C61A^ and 2000 SMARTA CD4 T cells x *Usp18*^C61A^ were mixed and transferred into WT mice. As control, same numbers of cell were transferred from P14 and SMARTA into WT mice. Mice were then infected with 2 x10^6^ PFU LCMV-Cl13 and survival was monitored (n = 5-6 mice/group).

**Supplementary Figure 6 (A)** *Usp18*^*fl/fl*^ and *Usp18*^*fl/fl*^ x CD11-Cre were infected with 2 x10^6^ PFU LCMV-Cl13. Survival was monitored (n = 6 mice/group). (B-D) *Usp18*^*fl/fl*^ and *Usp18*^*fl/fl*^ x CD11-Cre were infected with 2 x10^6^ PFU LCMV-Cl13, 8 days post infection, (B) total protein, LDH and IgM quantified by BCA and ELISA assays (C) indicated cytokines quantified by multiplex assay and (D) ISG15 were measured in the BALF by ELISA (n = 4-5 mice/group).

**Supplementary figure 7** (B) *Usp18*^*fl/fl*^ x LysM-Cre and WT mice were infected with 2 x10^6^ PFU LCMV-Cl13. 7 days post infection ISG15 was quantified in the BALF by ELISA (n = 4-7 mice/group). (B) *Usp18*^*fl/fl*^ x LysM-Cre and WT mice were infected with 2 x10^6^ PFU LCMV-Cl13. White blood cells (WBC) platelets and lymphocytes were measured in the blood at indicated time points (n = 8 mice/group).

**Supplementary figure 8** Gating strategy for sorting CD11b^+^ cells from the lung of WT and *Usp18*^C61A^ mice treated with anti-CD8 or isotype antibody on day -1 and 3 and infected with 2 x10^6^ PFU LCMV-Cl13.

**Supplementary figure 9** WT and *Usp18*^C61A^ mice were infected with Cl13 and treated with α-CD8 or isotype control. 7 days post infection, CD11b+ cells were purified from lung single cell suspension by flow cytometry, RNA isolated and sequenced using a ribo-depletion protocol. normalized counts of *Cxcr4* gene.

**Supplementary figure 10** MPO and MMP9 levels in BALF of WT, *Usp18*^C61A^ and *Usp18*^C61A^ treated with anti-CD8 or anti-IFN-γ or isotype antibody were quantified by ELISA assay.

## References

Alsohime, F., Martin-Fernandez, M., Temsah, M.H., Alabdulhafid, M., Le Voyer, T., Alghamdi, M., Qiu, X., Alotaibi, N., Alkahtani, A., Buta, S., et al. (2020). JAK Inhibitor Therapy in a Child with Inherited USP18 Deficiency. N Engl J Med 382, 256–265.

Andrews, S. (2010). A Quality Control Tool for High Throughput Sequence Data [Online].. (Available online at: http://www.bioinformatics.babraham.ac.uk/projects/fastqc/).

Arimoto, K.I., Lochte, S., Stoner, S.A., Burkart, C., Zhang, Y., Miyauchi, S., Wilmes, S., Fan, J.B., Heinisch, J.J., Li, Z., et al. (2017). STAT2 is an essential adaptor in USP18-mediated suppression of type I interferon signaling. Nat Struct Mol Biol 24, 279–289.

Baccala, R., Welch, M.J., Gonzalez-Quintial, R., Walsh, K.B., Teijaro, J.R., Nguyen, A., Ng, C.T., Sullivan, B.M., Zarpellon, A., Ruggeri, Z.M., et al. (2014). Type I interferon is a therapeutic target for virus-induced lethal vascular damage. Proc Natl Acad Sci U S A 111, 8925–8930.

Bogunovic, D., Byun, M., Durfee, L.A., Abhyankar, A., Sanal, O., Mansouri, D., Salem, S., Radovanovic, I., Grant, A.V., Adimi, P., et al. (2012). Mycobacterial disease and impaired IFN-gamma immunity in humans with inherited ISG15 deficiency. Science 337, 1684–1688.

Carevic, M., Singh, A., Rieber, N., Eickmeier, O., Griese, M., Hector, A., and Hartl, D. (2015). CXCR4+ granulocytes reflect fungal cystic fibrosis lung disease. Eur Respir J 46, 395–404.

Channappanavar, R., Fehr, A.R., Vijay, R., Mack, M., Zhao, J., Meyerholz, D.K., and Perlman, S. (2016). Dysregulated Type I Interferon and Inflammatory Monocyte-Macrophage Responses Cause Lethal Pneumonia in SARS-CoV-Infected Mice. Cell Host Microbe 19, 181–193.

Clausen, B.E., Burkhardt, C., Reith, W., Renkawitz, R., and Forster, I. (1999). Conditional gene targeting in macrophages and granulocytes using LysMcre mice. Transgenic Res 8, 265–277.

D’Cunha, J., Ramanujam, S., Wagner, R.J., Witt, P.L., Knight, E., Jr., and Borden, E.C. (1996). In vitro and in vivo secretion of human ISG15, an IFN-induced immunomodulatory cytokine. J Immunol 157, 4100–4108.

Dauphinee, S.M., Richer, E., Eva, M.M., McIntosh, F., Paquet, M., Dangoor, D., Burkart, C., Zhang, D.E., Gruenheid, S., Gros, P., et al. (2014). Contribution of increased ISG15, ISGylation and deregulated type I IFN signaling in Usp18 mutant mice during the course of bacterial infections. Genes Immun 15, 282–292.

Davidson, S., Crotta, S., McCabe, T.M., and Wack, A. (2014). Pathogenic potential of interferon alphabeta in acute influenza infection. Nat Commun 5, 3864.

Davidson, S., Maini, M.K., and Wack, A. (2015). Disease-promoting effects of type I interferons in viral, bacterial, and coinfections. J Interferon Cytokine Res 35, 252–264.

Ellis, T.N., and Beaman, B.L. (2004). Interferon-gamma activation of polymorphonuclear neutrophil function. Immunology 112, 2–12.

Evrard, M., Kwok, I.W.H., Chong, S.Z., Teng, K.W.W., Becht, E., Chen, J., Sieow, J.L., Penny, H.L., Ching, G.C., Devi, S., et al. (2018). Developmental Analysis of Bone Marrow Neutrophils Reveals Populations Specialized in Expansion, Trafficking, and Effector Functions. Immunity 48, 364–379 e368.

Gu, Z., Eils, R., and Schlesner, M. (2016). Complex heatmaps reveal patterns and correlations in multidimensional genomic data. Bioinformatics 32, 2847–2849.

Haegens, A., Vernooy, J.H., Heeringa, P., Mossman, B.T., and Wouters, E.F. (2008). Myeloperoxidase modulates lung epithelial responses to pro-inflammatory agents. Eur Respir J 31, 252–260.

Hartl, D., Krauss-Etschmann, S., Koller, B., Hordijk, P.L., Kuijpers, T.W., Hoffmann, F., Hector, A., Eber, E., Marcos, V., Bittmann, I., et al. (2008). Infiltrated neutrophils acquire novel chemokine receptor expression and chemokine responsiveness in chronic inflammatory lung diseases. J Immunol 181, 8053–8067.

Isles, H.M., Herman, K.D., Robertson, A.L., Loynes, C.A., Prince, L.R., Elks, P.M., and Renshaw, S.A. (2019). The CXCL12/CXCR4 Signaling Axis Retains Neutrophils at Inflammatory Sites in Zebrafish. Front Immunol 10, 1784.

Ketscher, L., Hannss, R., Morales, D.J., Basters, A., Guerra, S., Goldmann, T., Hausmann, A., Prinz, M., Naumann, R., Pekosz, A., et al. (2015). Selective inactivation of USP18 isopeptidase activity in vivo enhances ISG15 conjugation and viral resistance. Proc Natl Acad Sci U S A 112, 1577–1582.

Kim, J.V., Kang, S.S., Dustin, M.L., and McGavern, D.B. (2009). Myelomonocytic cell recruitment causes fatal CNS vascular injury during acute viral meningitis. Nature 457, 191–195.

Kim, K.I., Malakhova, O.A., Hoebe, K., Yan, M., Beutler, B., and Zhang, D.E. (2005). Enhanced antibacterial potential in UBP43-deficient mice against Salmonella typhimurium infection by up-regulating type I IFN signaling. J Immunol 175, 847–854.

Knight, E., Jr., and Cordova, B. (1991). IFN-induced 15-kDa protein is released from human lymphocytes and monocytes. J Immunol 146, 2280–2284.

Kolde, R. (2013). pheatmap: Pretty Heatmaps..

Lindner, H.A., Lytvyn, V., Qi, H., Lachance, P., Ziomek, E., and Menard, R. (2007). Selectivity in ISG15 and ubiquitin recognition by the SARS coronavirus papain-like protease. Arch Biochem Biophys 466, 8–14.

Liu, M., Hummer, B.T., Li, X., and Hassel, B.A. (2004). Camptothecin induces the ubiquitin-like protein, ISG15, and enhances ISG15 conjugation in response to interferon. J Interferon Cytokine Res 24, 647–654.

Love, M.I., Huber, W., and Anders, S. (2014). Moderated estimation of fold change and dispersion for RNA-seq data with DESeq2. Genome Biol. 15, 1–21.

Malakhov, M.P., Malakhova, O.A., Kim, K.I., Ritchie, K.J., and Zhang, D.E. (2002). UBP43 (USP18) specifically removes ISG15 from conjugated proteins. J Biol Chem 277, 9976–9981.

Malakhova, O., Malakhov, M., Hetherington, C., and Zhang, D.E. (2002). Lipopolysaccharide activates the expression of ISG15-specific protease UBP43 via interferon regulatory factor 3. J Biol Chem 277, 14703–14711.

Malakhova, O.A., Kim, K.I., Luo, J.K., Zou, W., Kumar, K.G., Fuchs, S.Y., Shuai, K., and Zhang, D.E. (2006). UBP43 is a novel regulator of interferon signaling independent of its ISG15 isopeptidase activity. EMBO J 25, 2358–2367.

McNab, F., Mayer-Barber, K., Sher, A., Wack, A., and O’Garra, A. (2015). Type I interferons in infectious disease. Nat Rev Immunol 15, 87–103.

Meuwissen, M.E., Schot, R., Buta, S., Oudesluijs, G., Tinschert, S., Speer, S.D., Li, Z., van Unen, L., Heijsman, D., Goldmann, T., et al. (2016). Human USP18 deficiency underlies type 1 interferonopathy leading to severe pseudo-TORCH syndrome. J Exp Med 213, 1163–1174.

Ng, C.T., Sullivan, B.M., Teijaro, J.R., Lee, A.M., Welch, M., Rice, S., Sheehan, K.C., Schreiber, R.D., and Oldstone, M.B. (2015). Blockade of interferon Beta, but not interferon alpha, signaling controls persistent viral infection. Cell Host Microbe 17, 653–661.

Nick, J.A., Caceres, S.M., Kret, J.E., Poch, K.R., Strand, M., Faino, A.V., Nichols, D.P., Saavedra, M.T., Taylor-Cousar, J.L., Geraci, M.W., et al. (2016). Extremes of Interferon-Stimulated Gene Expression Associate with Worse Outcomes in the Acute Respiratory Distress Syndrome. PLoS One 11, e0162490.

Oldstone, M.B. (2013). Lessons learned and concepts formed from study of the pathogenesis of the two negative-strand viruses lymphocytic choriomeningitis and influenza. Proc Natl Acad Sci U S A 110, 4180–4183.

Oldstone, M.B.A., Ware, B.C., Horton, L.E., Welch, M.J., Aiolfi, R., Zarpellon, A., Ruggeri, Z.M., and Sullivan, B.M. (2018). Lymphocytic choriomeningitis virus Clone 13 infection causes either persistence or acute death dependent on IFN-1, cytotoxic T lymphocytes (CTLs), and host genetics. Proc Natl Acad Sci U S A 115, E7814–E7823.

Owhashi, M., Taoka, Y., Ishii, K., Nakazawa, S., Uemura, H., and Kambara, H. (2003). Identification of a ubiquitin family protein as a novel neutrophil chemotactic factor. Biochem Biophys Res Commun 309, 533–539.

Perng, Y.C., and Lenschow, D.J. (2018). ISG15 in antiviral immunity and beyond. Nat Rev Microbiol 16, 423–439.

Radoshevich, L., Impens, F., Ribet, D., Quereda, J.J., Nam Tham, T., Nahori, M.A., Bierne, H., Dussurget, O., Pizarro-Cerda, J., Knobeloch, K.P., and Cossart, P. (2015). ISG15 counteracts Listeria monocytogenes infection. Elife 4.

Recht, M., Borden, E.C., and Knight, E., Jr. (1991). A human 15-kDa IFN-induced protein induces the secretion of IFN-gamma. J Immunol 147, 2617–2623.

Ritchie, K.J., Hahn, C.S., Kim, K.I., Yan, M., Rosario, D., Li, L., de la Torre, J.C., and Zhang, D.E. (2004). Role of ISG15 protease UBP43 (USP18) in innate immunity to viral infection. Nat Med 10, 1374–1378.

Schnell, F.J., Sundholm, S., Crumley, S., Iversen, P.L., and Mourich, D.V. (2012). Lymphocytic Choriomeningitis Virus Infection in FVB Mouse Produces Hemorrhagic Disease. Plos Pathog 8.

Schurmann, N., Forrer, P., Casse, O., Li, J., Felmy, B., Burgener, A.V., Ehrenfeuchter, N., Hardt, W.D., Recher, M., Hess, C., et al. (2017). Myeloperoxidase targets oxidative host attacks to Salmonella and prevents collateral tissue damage. Nat Microbiol 2, 16268.

Seiler, P., Aichele, P., Odermatt, B., Hengartner, H., Zinkernagel, R.M., and Schwendener, R.A. (1997). Crucial role of marginal zone macrophages and marginal zone metallophils in the clearance of lymphocytic choriomeningitis virus infection. Eur J Immunol 27, 2626–2633.

Shaabani, N., Honke, N., Nguyen, N., Huang, Z., Arimoto, K.I., Lazar, D., Loe, T.K., Lang, K.S., Prinz, M., Knobeloch, K.P., et al. (2018). The probacterial effect of type I interferon signaling requires its own negative regulator USP18. Sci Immunol 3.

Shaabani, N., Khairnar, V., Duhan, V., Zhou, F., Tur, R.F., Haussinger, D., Recher, M., Tumanov, A.V., Hardt, C., Pinschewer, D., et al. (2016). Two separate mechanisms of enforced viral replication balance innate and adaptive immune activation. J Autoimmun 67, 82–89.

Speer, S.D., Li, Z., Buta, S., Payelle-Brogard, B., Qian, L., Vigant, F., Rubino, E., Gardner, T.J., Wedeking, T., Hermann, M., et al. (2016). ISG15 deficiency and increased viral resistance in humans but not mice. Nat Commun 7, 11496.

Street, K., Risso, D., Fletcher, R.B., Das, D., Ngai, J., Yosef, N., Purdom, E., and Dudoit, S. (2018). Slingshot: cell lineage and pseudotime inference for single-cell transcriptomics. BMC Genomics 19, 477.

Stuart, T., Butler, A., Hoffman, P., Hafemeister, C., Papalexi, E., Mauck, W.M., III, Hao, Y., Stoeckius, M., Smibert, P., and Satija, R. (2019). Comprehensive Integration of Single-Cell Data. Cell 177, 1888-1902.e1821.

Sunderkotter, C., Nikolic, T., Dillon, M.J., Van Rooijen, N., Stehling, M., Drevets, D.A., and Leenen, P.J. (2004). Subpopulations of mouse blood monocytes differ in maturation stage and inflammatory response. J Immunol 172, 4410–4417.

Swaim, C.D., Scott, A.F., Canadeo, L.A., and Huibregtse, J.M. (2017). Extracellular ISG15 Signals Cytokine Secretion through the LFA-1 Integrin Receptor. Mol Cell 68, 581–590 e585.

Team, R.D.C. (2008). R: A language and environment for statistical computing. R Foundation for Statistical Computing. (Vienna, Austria).

Teijaro, J.R. (2015). The role of cytokine responses during influenza virus pathogenesis and potential therapeutic options. Curr Top Microbiol Immunol 386, 3–22.

Teijaro, J.R. (2016). Type I interferons in viral control and immune regulation. Curr Opin Virol 16, 31–40.

Teijaro, J.R., Ng, C., Lee, A.M., Sullivan, B.M., Sheehan, K.C., Welch, M., Schreiber, R.D., de la Torre, J.C., and Oldstone, M.B. (2013). Persistent LCMV infection is controlled by blockade of type I interferon signaling. Science 340, 207–211.

Teijaro, J.R., Walsh, K.B., Cahalan, S., Fremgen, D.M., Roberts, E., Scott, F., Martinborough, E., Peach, R., Oldstone, M.B., and Rosen, H. (2011). Endothelial cells are central orchestrators of cytokine amplification during influenza virus infection. Cell 146, 980–991.

Walsh, K.B., Teijaro, J.R., Wilker, P.R., Jatzek, A., Fremgen, D.M., Das, S.C., Watanabe, T., Hatta, M., Shinya, K., Suresh, M., et al. (2011). Suppression of cytokine storm with a sphingosine analog provides protection against pathogenic influenza virus. Proc Natl Acad Sci U S A 108, 12018–12023.

Wilson, E.B., Yamada, D.H., Elsaesser, H., Herskovitz, J., Deng, J., Cheng, G., Aronow, B.J., Karp, C.L., and Brooks, D.G. (2013). Blockade of chronic type I interferon signaling to control persistent LCMV infection. Science 340, 202–207.

Yamada, M., Kubo, H., Kobayashi, S., Ishizawa, K., He, M., Suzuki, T., Fujino, N., Kunishima, H., Hatta, M., Nishimaki, K., et al. (2011). The increase in surface CXCR4 expression on lung extravascular neutrophils and its effects on neutrophils during endotoxin-induced lung injury. Cell Mol Immunol 8, 305–314.

Yoshida, H., Lareau, C.A., Ramirez, R.N., Rose, S.A., Maier, B., Wroblewska, A., Desland, F., Chudnovskiy, A., Mortha, A., Dominguez, C., et al. (2019). The cis-Regulatory Atlas of the Mouse Immune System. Cell 176, 897-912.e820.

