## Supplementary figures and images for "ISG15 drives immune pathology and respiratory failure during viral infection"

### Supplementary Figure 1

#### Supplementary Figure 1

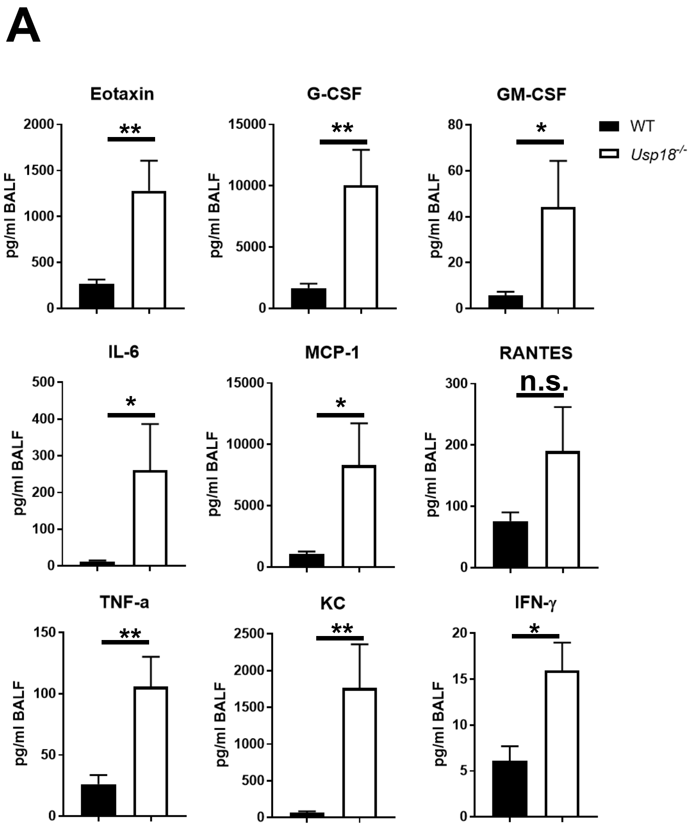


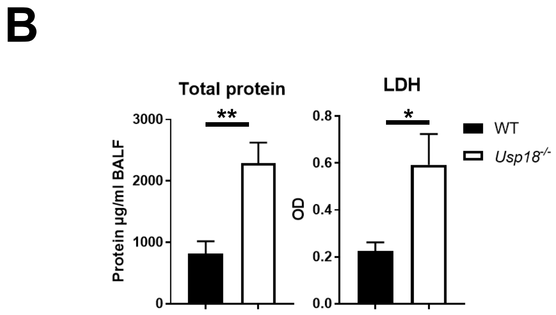

### Supplementary Figure 2

#### Supplementary Figure 2

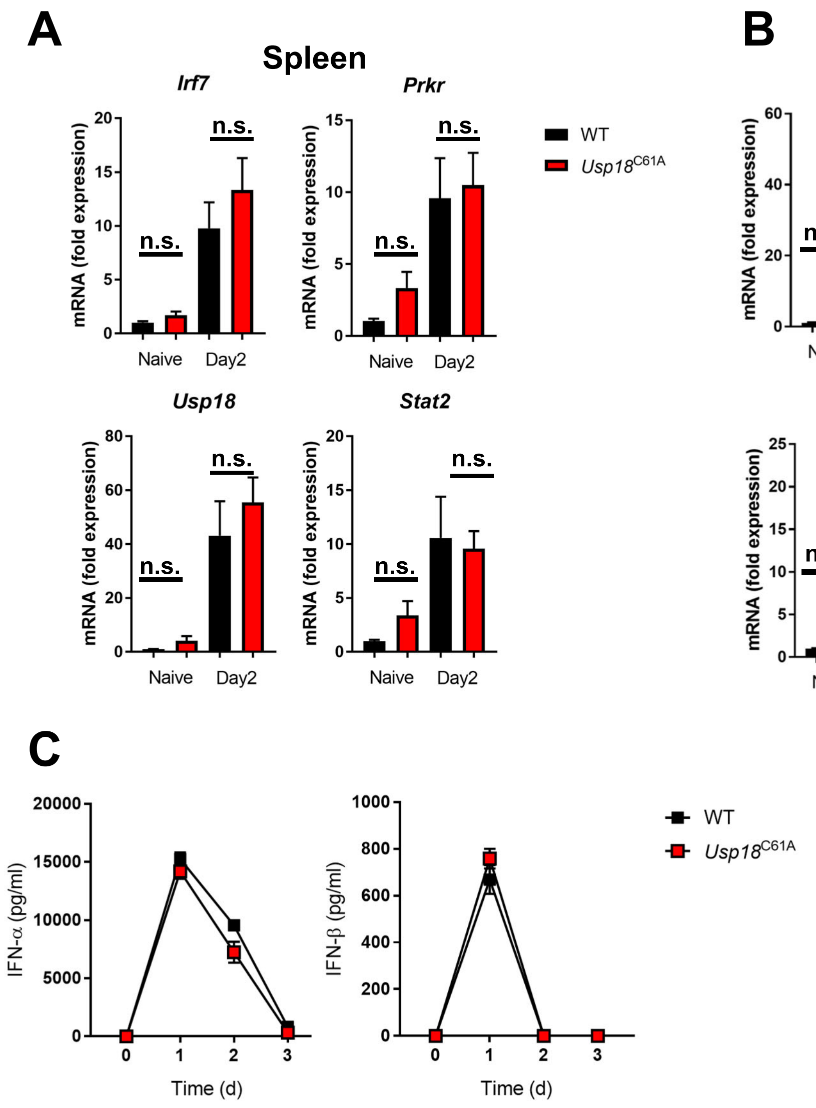

### Supplementary Figure 3

#### Supplementary Figure 3

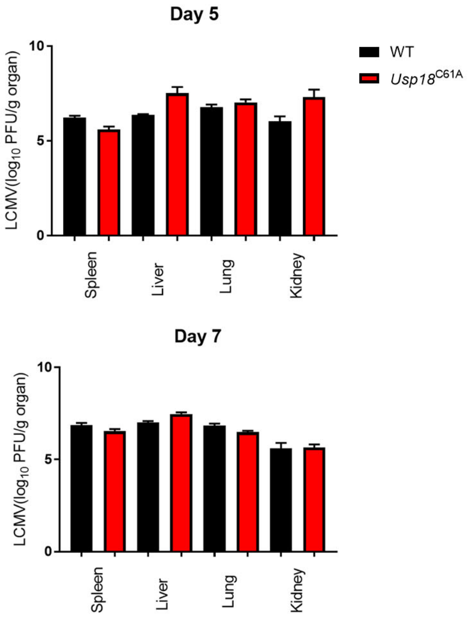

### Supplementary Figure 4

#### Supplementary Figure 4

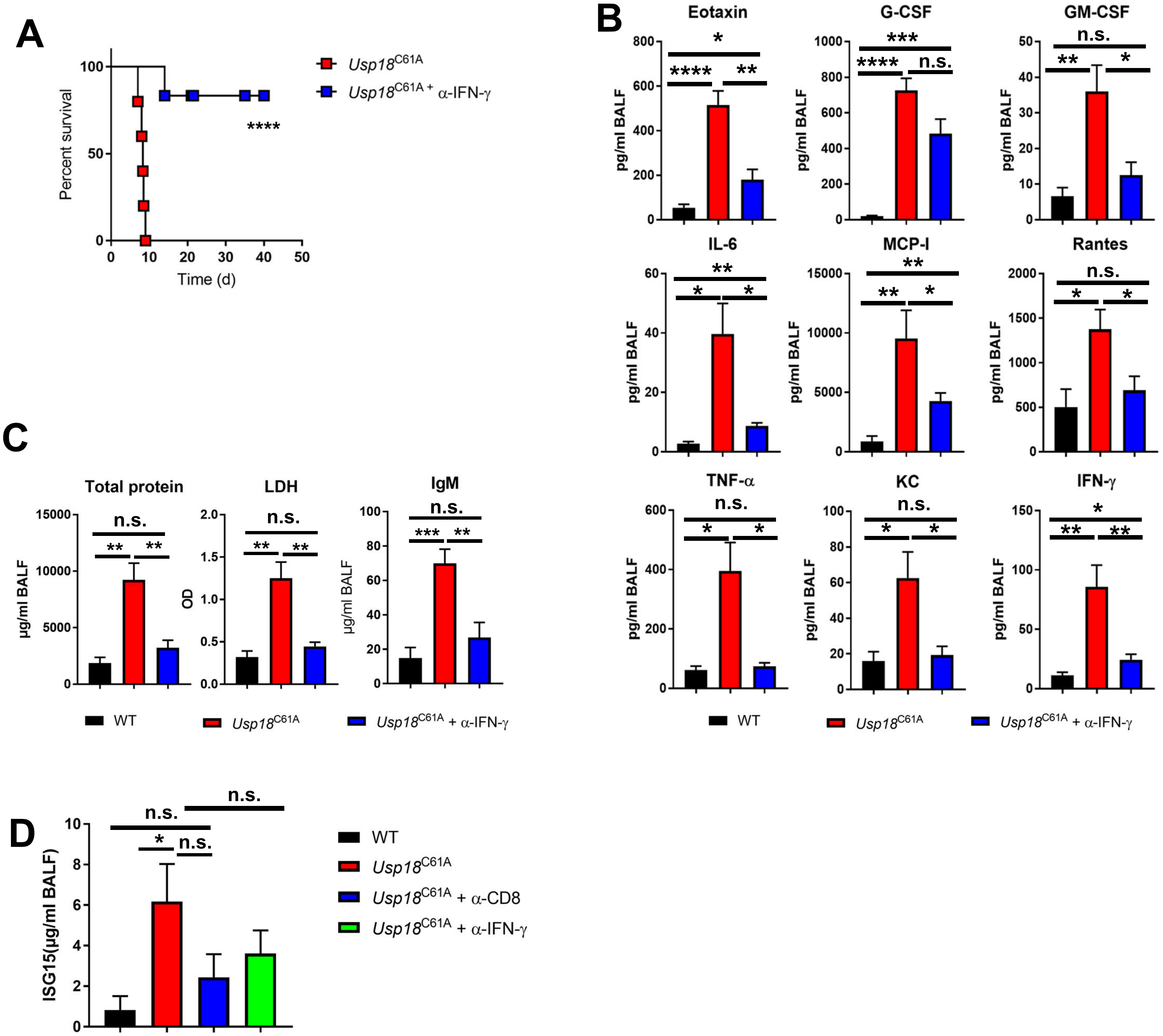

### Supplementary Figure 5

#### Supplementary Figure 5

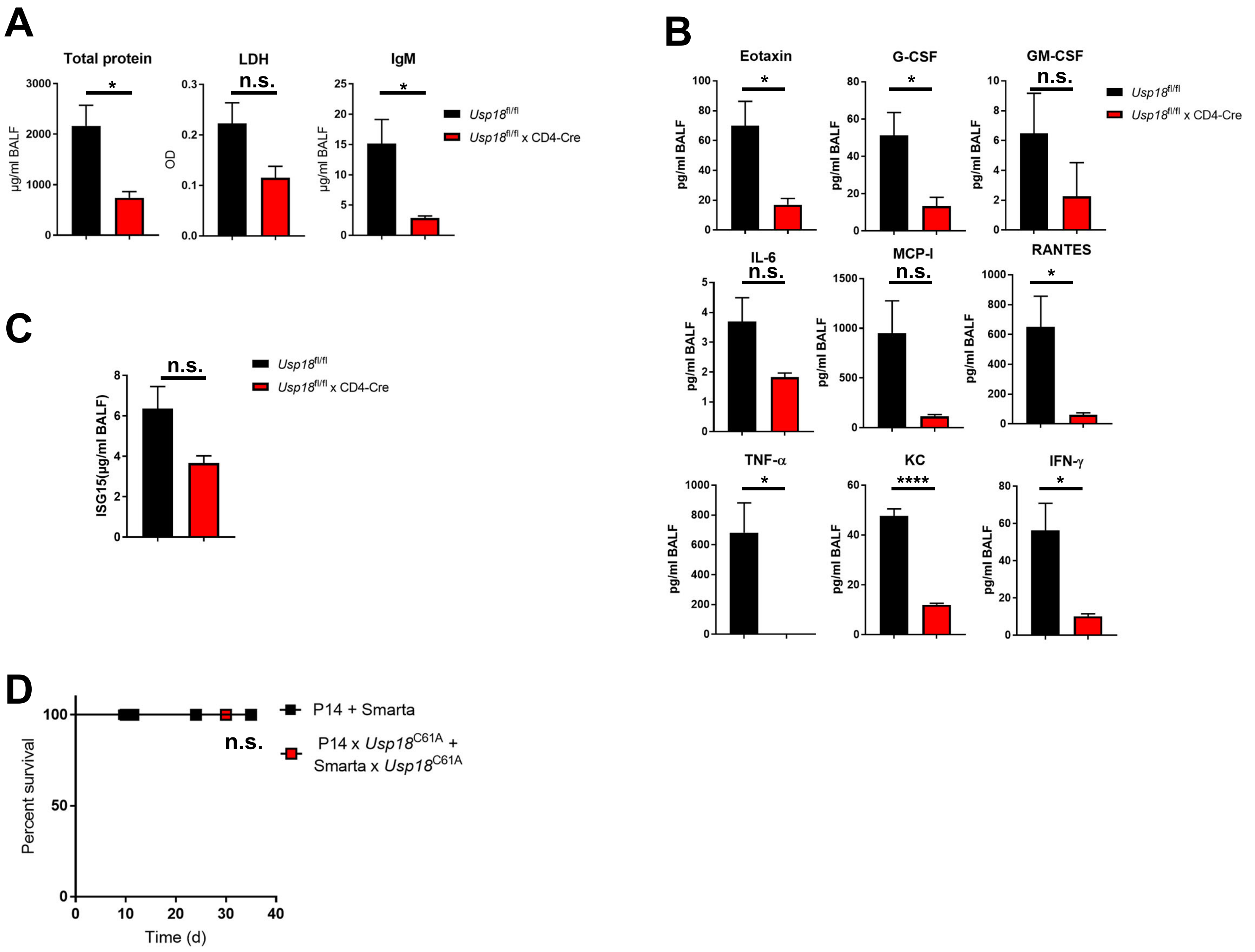

### Supplementary Figure 6

#### Supplementary Figure 6

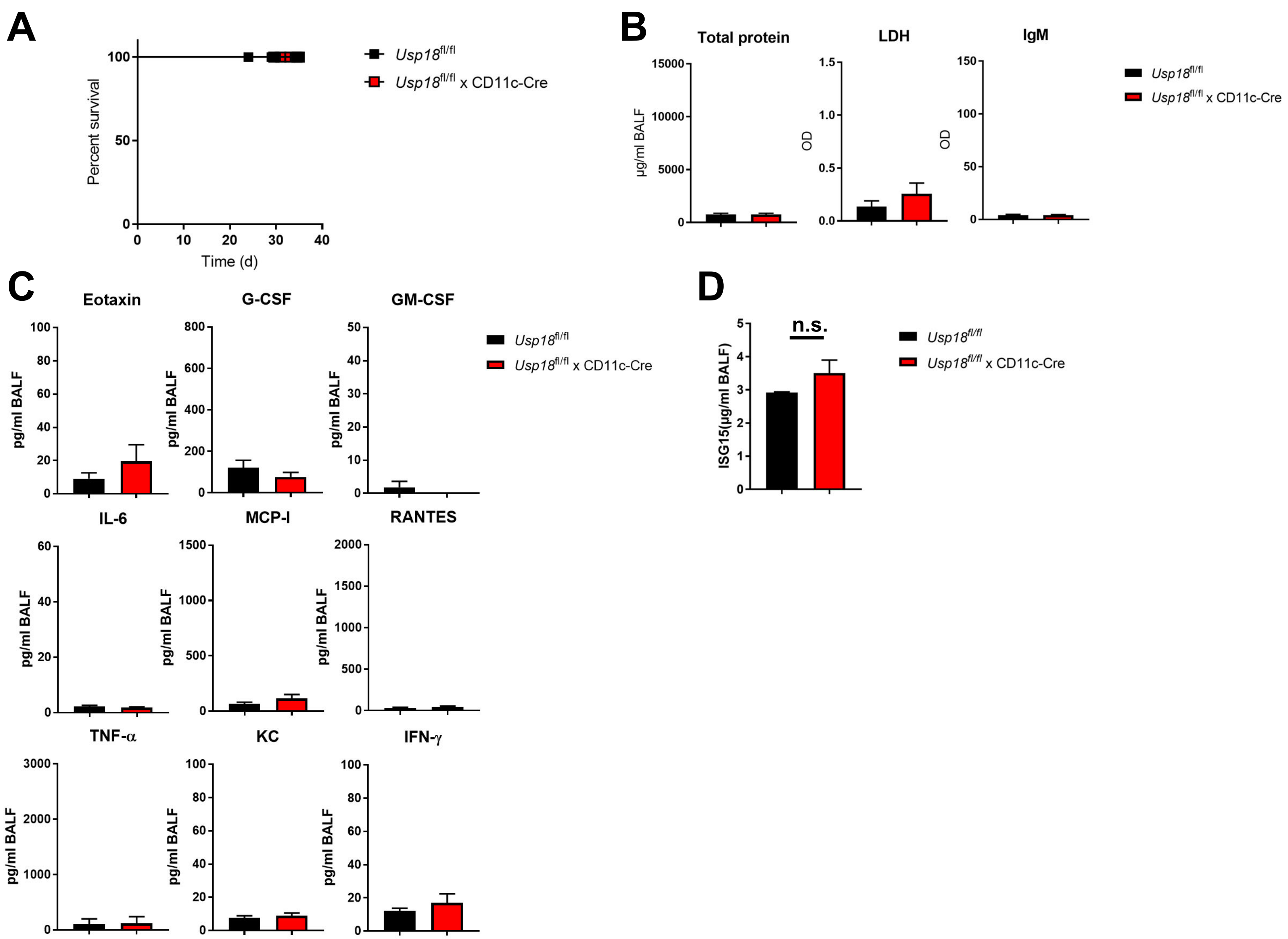

### Supplementary Figure 7

#### Supplementary Figure 7

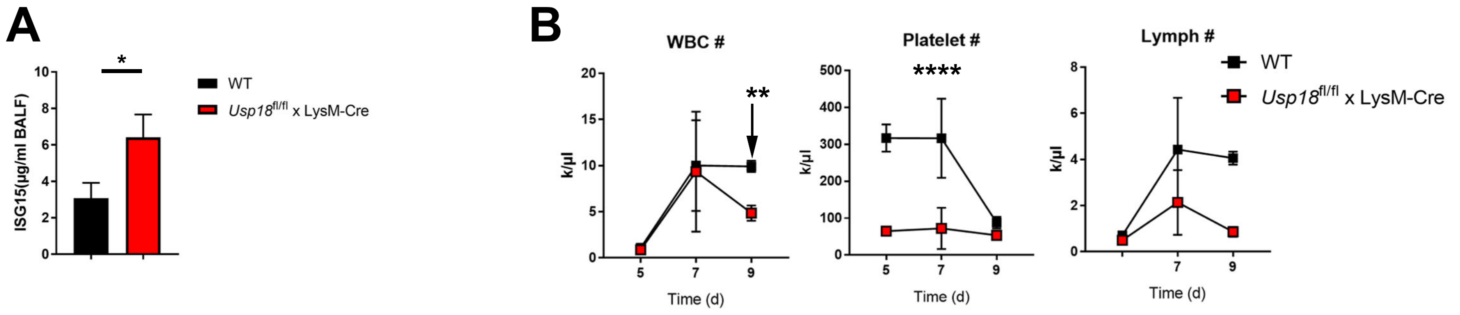

### Supplementary Figure 8

#### Supplementary Figure 8

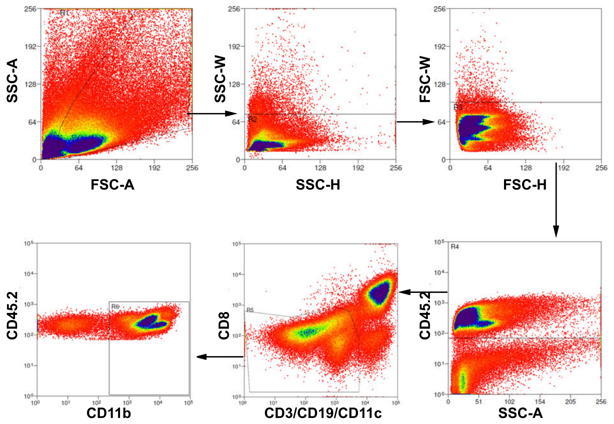

### Supplementary Figure 9

#### Supplementary Figure 9

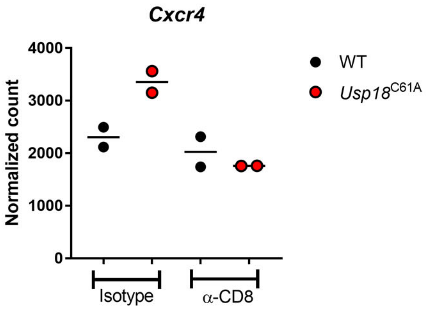

### Supplementary Figure 10

#### Supplementary Figure 10

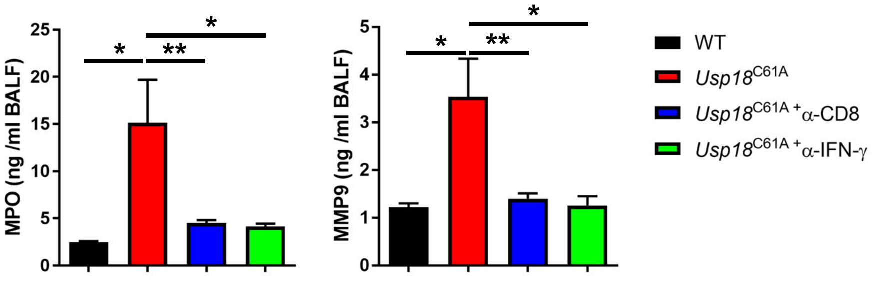
